# GLOBAL HOST RESPONSES TO THE MICROBIOTA AT SINGLE CELL RESOLUTION IN GNOTOBIOTIC ZEBRAFISH

**DOI:** 10.1101/2022.03.28.486083

**Authors:** Michelle S. Massaquoi, Garth Kong, Daisy Chilin, Mary K. Hamilton, Ellie Melancon, Judith S. Eisen, Karen Guillemin

**Affiliations:** Institute of Molecular Biology, University of Oregon, 1318 Franklin Blvd, Eugene, OR 97403, USA; Institute of Neuroscience, University of Oregon, 1254 University of Oregon, Eugene, OR 97403; Humans and the Microbiome Program, CIFAR, Toronto, ON M5G 1M1, Canada

**Keywords:** microbiota, zebrafish, development, transcriptomics, gnotobiology

## Abstract

Resident microbes are a feature of vertebrate animals that influence diverse aspects of their biology from tissue development to metabolism. Here we describe transcriptional responses to the microbiota across all the cells of a model developing vertebrate, the larval zebrafish. By performing single cell transcriptomic analysis of whole germ free and conventionalized larvae, we show that the impacts of the microbiota are sensed by all major organ systems but that responses are highly specific to different cell types. The presence of microbiota stimulates the expansion of progenitor-like cells in epithelial tissues and increases proliferation gene expression in progenitor-like cell populations of the immune and nervous systems. Across many cell types, including enterocytes, immune cells, and neurons, the microbiota upregulates expression of genes involved in microbial responses, cell type-specific activities, and cell type-specific deployment of ATP metabolism genes. These combined transcriptional patterns demonstrate how the microbiota simultaneously modulate cellular immune and metabolic programs. The impacts of the microbiota on tissue development are illustrated by the exocrine pancreas, which in the absence of the microbiota is smaller and composed of uniformly differentiated acinar cells. The presence of the microbiota results in exocrine pancreas enlargement and heterogeneous cellular expression of digestive enzyme and secretion genes, demonstrating how the microbiota promotes plasticity in tissue development and function. This single cell transcriptional dataset demonstrates the impacts of the microbiota on vertebrate development across the body and provides a foundation for dissecting cell type specific responses to microbial consortia members or molecules.

**Summary:** Animal development proceeds in the presence of intimate microbial associations, but the extent to which different host cells across the body respond to resident microbes remains to be fully explored. Using the vertebrate model organism, the larval zebrafish, we assessed transcriptional responses to the microbiota across the entire body at single cell resolution. We find that cell types across the body, not limited to tissues at host-microbe interfaces, respond to the microbiota. Responses are cell-type specific, but across many tissues the microbiota enhances cell proliferation, increases metabolism, and stimulates a diversity of cellular activities, revealing roles for the microbiota in promoting developmental plasticity. This work provides a resource for exploring transcriptional responses to the microbiota across all cell types of the vertebrate body and generating new hypotheses about the interactions between vertebrate hosts and their microbiota.

## Introduction

Bacteria were the first organisms to evolve on earth, preceding the emergence of animals by roughly 3 billion years. All animals evolved amidst the pressures exerted by the abundant bacteria, fungi and viruses in their world, which necessitated cellular and tissue strategies to co-exist with resident microbial communities, or microbiota, that live on and within animal bodies. Emerging insights into the many roles that microbiotas play in host development and homeostasis are redefining how we view animal biology (Mcfall-Ngai, 2014). The study of animal-microbiota interactions are being transformed by new technologies for single cell characterizations (Sharma and Thaiss, 2020), but the impacts of resident microbes on cellular transcription have not yet been described globally across an entire vertebrate animal body. The goal of this research is to create a resource that catalogues cell type specific transcriptional responses of the larval zebrafish to its microbiota. Zebrafish are an excellent model in which to study impacts of the microbiota on vertebrate biology because they are readily amendable to gnotobiotic manipulations (Melancon *et al*., 2017), their small size allows single cell transcriptional profiling of the entire body (Farnsworth, Saunders and Miller, 2019), and they share common developmental and physiological programs with larger vertebrates such as humans. In this study, we used the 10x Genomics pipeline to interrogate the transcriptomes of single cells dissociated from cohorts of larval zebrafish reared in the presence or absence of their normal microbiota. We report our analysis of the global, tissue, and cell type responses to the microbiota that emerge from this dataset, which will serve as a valuable hypothesis generating resource for the microbiome sciences field.

## Results & Discussion

### Responsiveness to microbes is widespread across cells of the vertebrate body

The small size and tractable gnotobiology of zebrafish larvae afford an unprecedented opportunity to survey the responsiveness of every vertebrate host cell type to the presence of their microbiota. We derived sterile zebrafish embryos and reared them germ-free (GF) or conventionalized them (CVZ) them with parental tank water. At 6 days post fertilization (dpf) whole larvae were dissociated into single cells and processed for single cell RNA sequencing (Fig. 1A). This analysis yielded 33, 992 cells that averaged 23,485 mean reads/cell, with 879 median genes/cell, 2,396 unique transcript molecules (UMI)/cell and 22,953 and 22,898 total genes detected from CVZ and GF groups, respectively.

**Figure 1.**
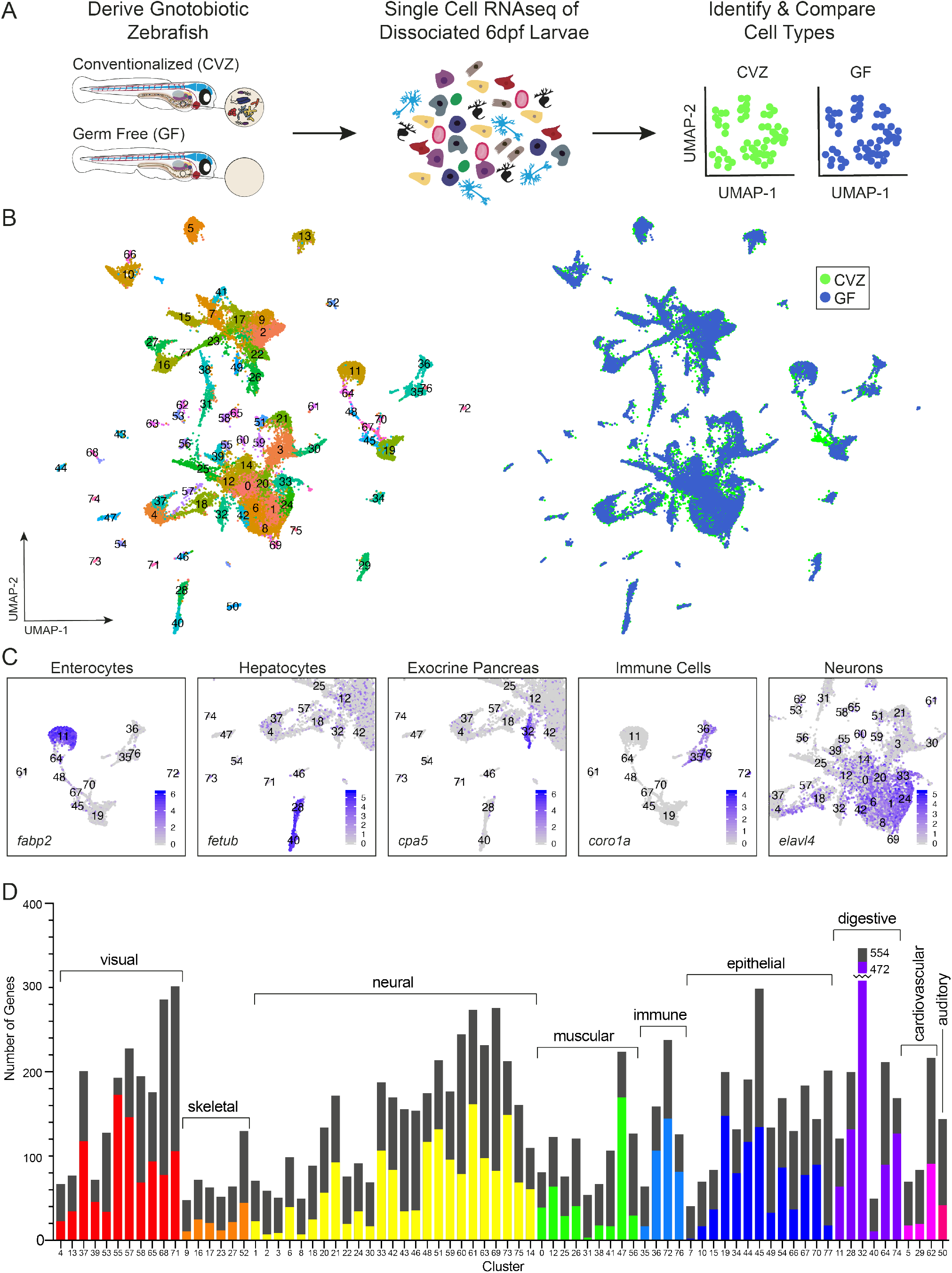
Single cell transcriptional analysis of whole gnotobiotic larval zebrafish. A) Whole conventionalized (CVZ) and germ free (GF) 6dpf zebrafish larvae were dissociated into individual cells prior to single cell RNA sequencing. B) uMAP plots display 78 cell type clusters shared between experimental groups. C) Cell types were identified by their transcriptomic profile; examples (with marker genes) include enterocytes (*fabp2*), hepatocytes (*fetub*), acinar cells of the exocrine pancreas (*cpa5*), immune cells (*coro1a*) and neurons (*elavl4*). D) Host cell types differentially respond to the presence of the microbiota throughout the body as illustrated by the total number of genes significantly enriched within each experimental group (p<0.05). Bars represent the sum of differentially expressed genes in CVZ and GF cells for each cluster with the colored portion representing the number of genes significantly more highly expressed within CVZ versus GF cells and the black portion representing the number of genes significantly more highly expressed in GF versus CVZ cells.

Based on transcriptional profiles, our dataset is composed of diverse host cell types with largely equal representation from CVZ and GF groups (Fig. 1B). We used the Seurat package in R to integrate cells from both treatments (Stuart *et al*., 2018), identify common sources of variation between CVZ and GF cells, and perform a linear reduction of the data by principal component analysis (PCA). In the integration strategy, matching cells from each dataset are anchored together based on the similarity of their transcriptional profile. Within integrated datasets, clusters that include similar numbers of cells from each treatment group represent the pairing of cells in a similar biological state. The JackStrawPlot, ElbowPlot and HeatMap functions included in Seurat deciphered which principal components (PC) represent a robust compression of the data (SFig. 1&2). To confirm clustering of transcriptionally similar cells across experimental groups by the Seurat integration strategy, we aligned 6dpf CVZ cells with different developmental timepoints (1, 2, 5 dpf) of whole organism dissociations from the Zebrafish Single Cell Atlas (Farnsworth, Saunders and Miller, 2019)(SFig. 3A&B). As anticipated, cells from our 6dpf CVZ zebrafish aligned more consistently with 5dpf cells of the atlas compared to 1 and 2dpf cells.

By integrating CVZ and GF cells together, we generated lists of significantly enriched genes shared by CVZ and GF cells for each cluster, allowing identification of different cell types present in both treatment groups. Cell type identities were assigned based on tissue location annotations from the Zebrafish Information Network (ZFIN) (Howe *et al*., 2013) of the top enriched genes for each cluster and annotations from the Zebrafish Single Cell Atlas (S.Table1) (Fig. 1C). This annotation established that the dataset contains diverse cell types from all major organ systems, enabling us to compare transcriptional differences between CVZ and GF treatments for cells across the vertebrate body.

Comparing differentially represented transcripts between CVZ and GF cells for each cluster revealed host cell responses to the microbiota occur across many cell types, not limited to cells in direct contact with microbes (Fig. 1D). Additionally, the high variability in the number of enriched genes across cell types demonstrates that different cells respond differently to the microbiota even within a tissue type. One cluster, number 45, lacked GF cell representation (Fig. 1B) in a pattern that was robust to different permutations of PC number (SFig. 4A). This cluster is enriched for genes typically found within epithelia, such as *epcam*, and gene ontology (GO) analysis showed a signature of tissue regeneration and immune activation. Although some of the cells express markers of skin epithelia, such as different types of *keratins*, the cluster lacks a strong signature of tissue identity (SFig. 4B-D). Based on this gene expression pattern and further analysis of dissected intestines discussed below, this cluster appears to consist of progenitor cells from multiple epithelial tissues that co-cluster based on their common cellular responses to microbiota associated with tissue growth and regeneration programs. The absence of this cell population in the GF dataset is consistent with the microbiota’s role in stimulating epithelial cell proliferation in tissues including intestine (Rawls, Samuel and Gordon, 2004; Cheesman *et al*., 2011) and skin (Meisel et al., 2018). Together, these analyses illustrate that cells across the vertebrate body are responsive to the presence of the microbiota, which promotes tissue development. Below we discuss cell type specific responses to the microbiota.

### The microbiota stimulates intestinal epithelial cell differentiation and function

Transcriptional responses to the microbiota have been best characterized within the intestine (reviewed in (Heppert *et al*., 2021)). To supplement our whole larval single cell analysis and compare it to previous analyses, we conducted an additional experiment in which we dissected larval digestive systems, comprised of intestines and associated liver and pancreas tissue, dissociated the cells, and performed single cell sequencing (SFig. 5A&B). Data from dissected digestive tracts validated our annotation of intestinal epithelium, liver, and pancreas cells in the whole larva dataset (Fig. 2A&B). Comparing our lists of enriched transcripts in digestive tract clusters with previous microarray (Rawls, Samuel and Gordon, 2004) and single cell RNAseq (Willms *et al*., 2022) analysis of dissected CVZ and GF larval zebrafish digestive tracts revealed good concordance between these datasets (STable. 2&3; SFig. 6).

**Figure 2.**
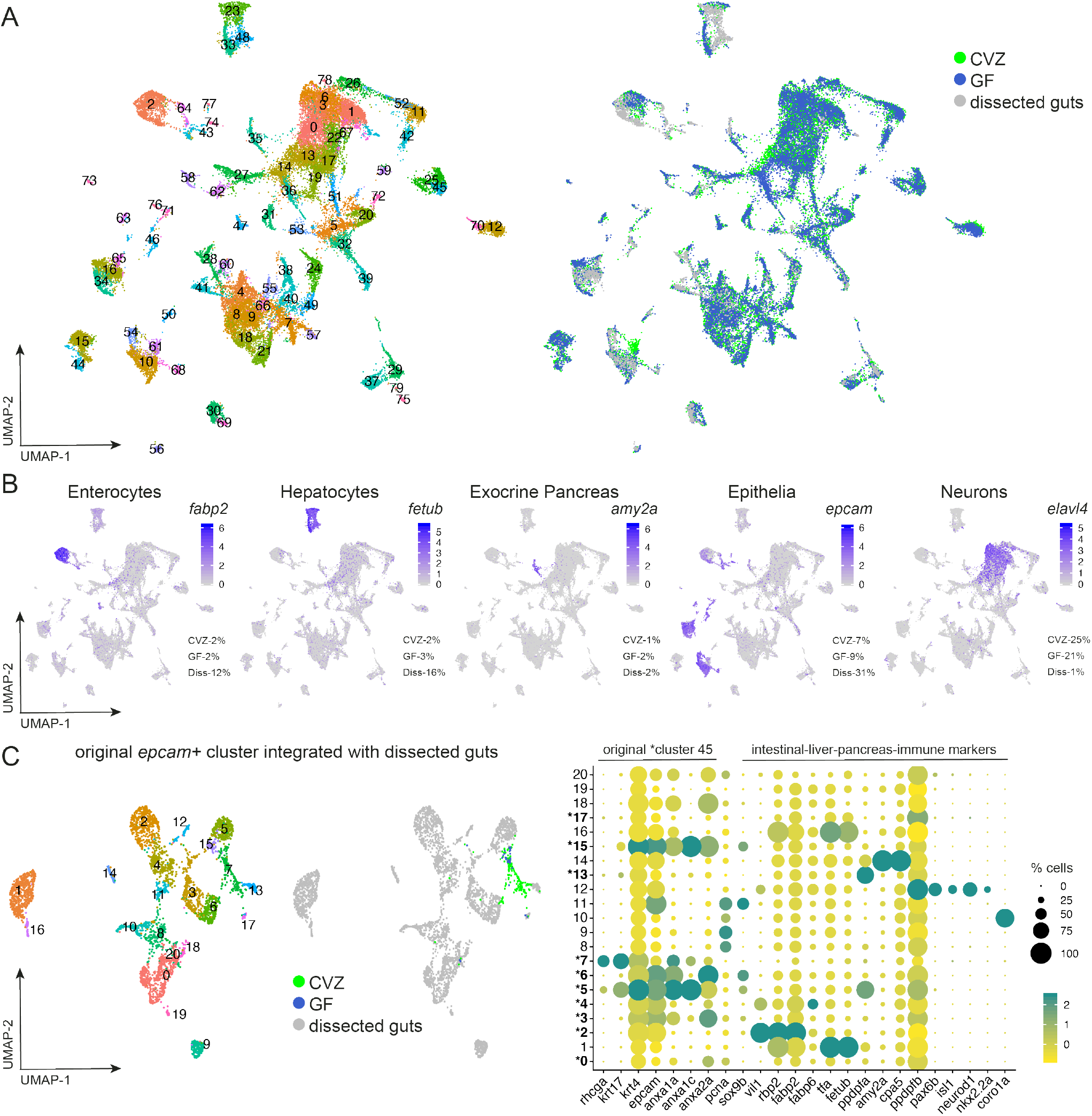
Integration of whole gnotobiotic larval zebrafish cells with dissected larval digestive system cells. A) uMAP plots display integration of single cells dissociated from whole larvae and dissected larval guts. B) Clusters populated by cells from each experimental group confirm and illustrate digestive cell types from dissociation of whole larvae including enterocytes (*fabp2*), hepatocytes (*fetub*), acinar cells of the exocrine pancreas (*amy2a*), and epithelial cells (*epcam*). The percentage of cells within designated clusters, illuminated by different biomarkers, with respect to total cells for each experimental group are shown. Cells from dissected digestive systems primarily populated clusters from whole larvae annotated as digestive system cell types and were a small percentage of the neurons(*elavl4*) cluster, verifying the integration strategy. C) uMAP plots show that cells from original epithelial cluster 45 include a minority of cells likely to be of the digestive system. The dotplot, and all following dotplots included in the manuscript, displays both the percentage of cells within a cluster or subcluster expressing a transcript (dot diameter) and expression level of the transcript (dot color). Expression of genes within this dotplot for a subcluster is relative to all other subclusters. Bolded subcluster numbers designated with an * represent subclusters that include cell(s) from original cluster 45.

### Progenitor cells in the CVZ dataset include digestive system cells

Upon integrating the dissected digestive system cells with the whole organism dataset, we observed that CVZ epithelial cells from original cluster 45 maintained their co-segregation into a cluster assigned as 61 but containing fewer cells than the original 45 (Fig. 2A&B). We hypothesized that cells from the original cluster 45 had clustered with digestive cell types enriched in the dissected digestive system dataset. To investigate this possibility, we computationally isolated cluster 45 cells and used Seurat to integrate them with the dissected digestive system data (Fig. 2C). A majority of CVZ cells segregated to subcluster 7, indicating that most cells from original cluster 45 are extra-intestinal, likely including skin cells. However, 12% of the CVZ cells spread to diverse digestive system subclusters, suggesting that original cluster 45 includes progenitors of different digestive system tissues.

### Intestinal enterocytes segregate by region and function

We identified two adjacent clusters of intestinal enterocytes based on regional specific genes (defined by (Lickwar *et al*., 2017)) including the proximal intestine genes *rbp2a* and *chia*.*2* (cluster 11, Fig. 3A) and the ileum-specific *fabp6* (cluster 64, SFig. 7A). To further delineate cell types within clusters 11 and 64, we isolated and re-clustered these data (SFig. 7B&C), which resolved the proximal, mid, and distal intestine described in (Lickwar *et al*., 2017) and (Wen *et al*., 2021), and was consistent with further analysis of the Zebrafish Atlas intestinal epithelia clusters (Postlethwait et al., 2020). GO analysis of significantly enriched genes within cluster 11 of CVZ versus GF enterocytes revealed that the microbiota stimulates specific cellular responses. We observed enrichment of genes involved in macromolecule catabolism, metallopeptidase activity, and lipid-protein assembly, including *apoa4b*.*1* and *apoa4b*.*2*, consistent with previous work illustrating that the microbiota regulates fatty acid metabolism and intestinal absorption (Semova *et al*., 2012). We also saw induction of responses to microbes such as chitinase genes *chia*.*1* and *chia*.*2*, and other cellular responses consistent with microbial challenge including responses to inorganic compounds, temperature, and ER stress (Fig. 3A). Enrichment of *chia*.*1, muc13b and tmem1761*.*2* within CVZ versus GF enterocytes was also observed in an analysis of differential gene expression between CVZ and GF 6dpf dissected digestive systems (Willms et al. 2022). Consistent with our annotation of cluster 64 as mid intestine enterocytes, which includes lysosome rich enterocytes with high capacity for absorption and processing of luminal proteins (Park *et al*., 2019), we observed significant enrichment within CVZ cells of genes involved in coated vesicle and proteosome complex formation.

**Figure 3.**
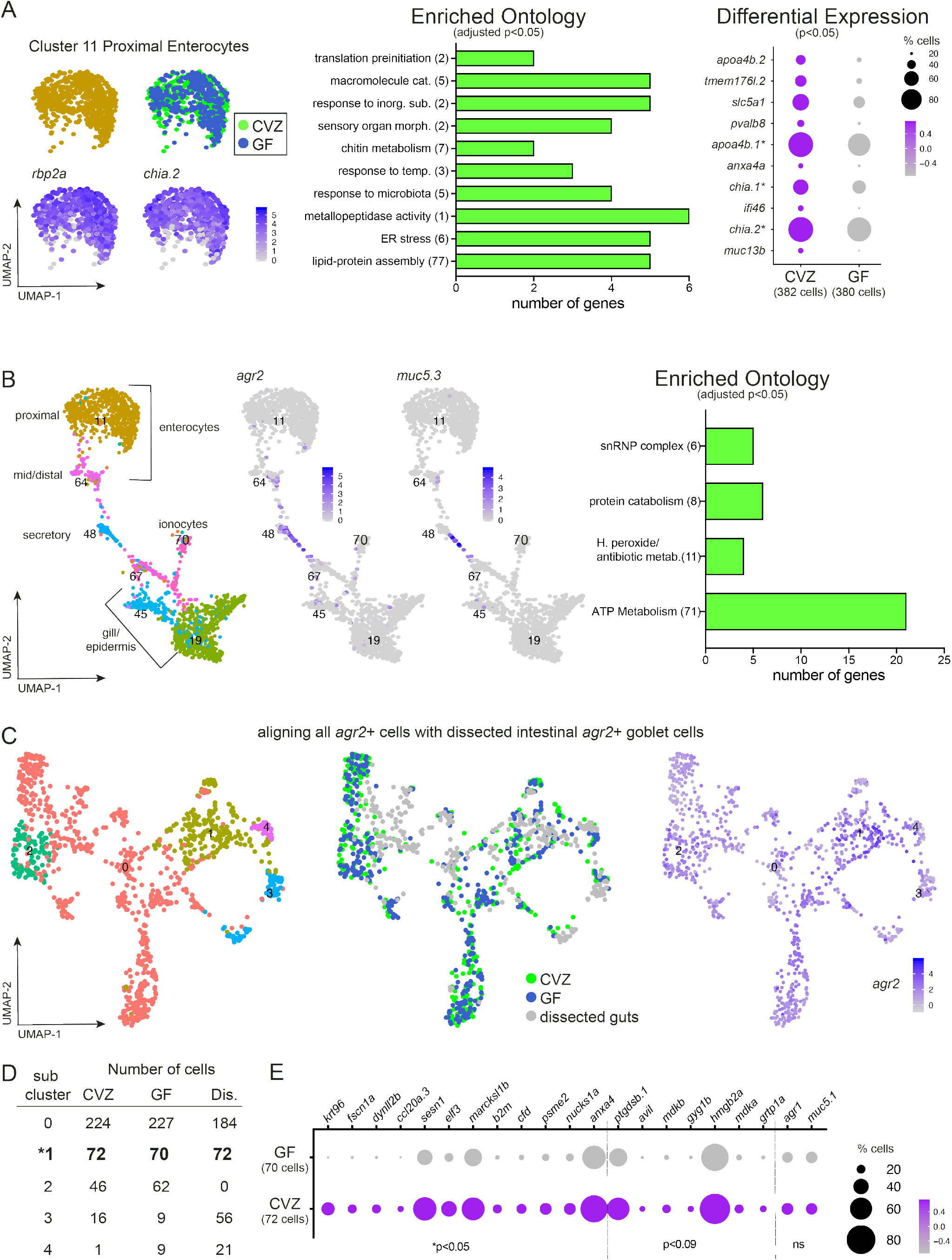
Intestinal enterocytes and secretory cells exhibit cell type specific transcriptional responses to the microbiota. A) Cluster 11 is composed of enterocytes from the proximal intestine marked by high expression of *rbp2a* and *chia*.*2*. GO analysis plot represents gene function categories enriched in CVZ versus GF cells in cluster 11 and the dotplot shows genes significantly enriched within CVZ versus GF cells of cluster 11. Genes included in the dotplot designated with an * correspond to genes included in GO categories. B) Mucin-secreting cells are localized to cluster 48 indicated by expression of *agr2* and *muc5*.*3*. GO analysis plot is based on genes significantly enriched within CVZ versus GF cells of cluster 48. C) uMAP plots display integration of all *agr2*+ cells from whole larvae with *agr2*+ cells from dissected guts. By uMAP and feature plots, subcluster1 displays the most overlap of cells from each experimental group and the highest expression of *agr2*. D) Subcluster 1 demonstrates consistent pairing of cells from each experimental group from the integration. E) Dotplot illustrates enriched expression of several genes within CVZ cells of subcluster 1. GO plots in this figure and all following figures use bars to show the total number of genes included in a GO category. The number in parentheses displayed after a GO category title on the y-axis signifies the number of GO terms binned into the category. GO terms are considered significant at p<0.05 after an fdr p-corrected adjustment.

### Intestinal goblet cells segregate with other mucus secreting cells

Goblet cells within the intestinal epithelium secrete mucins into the lumen, providing a nutrient source for microbes as well as a mucus barrier between the luminal microbiota and the epithelium. Agr2 is an endoplasmic reticulum protein disulfide-isomerase that is highly expressed in mucus secretory cells including goblet cells of the zebrafish intestine, pharynx and epidermis (Shih *et al*., 2007). Clusters 50 and 48 showed the highest enrichment for *agr2* (~60% and ~30%, *agr2*+ cells, respectively). To identify intestinal goblet cells within the *agr2*+ cell populations, we looked for expression of zebrafish orthologs of *Muc2* and *Muc5*, mucins specific to mouse intestinal goblet cells (Larsson et al. 2011). We did not observe *muc2*.*2* expression in our data set of whole or dissected digestive systems or in the Zebrafish Atlas, however *muc5*.*1* and *muc5*.*3* expression were enriched within cluster 48, which lies between the intestinal and epidermal epithelium clusters (S.Table1) (Fig. 3B). GO analysis showed significant enrichment of genes involved in hydrogen peroxide and antibiotic metabolism within CVZ cells of cluster 48 relative to GF, consistent with protective responses elicited in cells in close proximity to microbes. Because expression of *muc5*.*3* in zebrafish has been found within the esophagus and intestine (Jevtov *et al*., 2014; Udayangani *et al*., 2017), we speculate that cluster 48 includes extra-intestinal goblet cells.

To refine our identification of intestinal goblet cells, we analyzed *agr2* expression in the dissected digestive system dataset (SFig. 5B). The *agr2*+ population within cluster 4 was enriched for intestinal epithelial markers, suggesting that these are intestinal secretory cells (SFig. 5B). Using the Seurat integration method, we combined all CVZ and GF *agr2*+ cells from whole larvae with the dissected intestinal *agr2*+ population from cluster 4 (Fig. 3C; SFig. 5B). In this analysis, subcluster 1 exhibited the tightest co-clustering of cells from both datasets, suggesting these *agr*+ cells are intestinal goblet cells (Fig. 3C&D). Compared to GF cells within subcluster 1, CVZ cells had a significant enrichment of genes involved in cytoskeleton architecture (*krt96, fscn1a, dynll2b*), immune responses (*ccl20a*.*3, sesn1, b2m*), and transcription factor activity and development, including Notch signaling (*elf3, marcksl1b, cfd, psme2, nucks1a*) (Fig. 3E). Although not statistically significant, several genes involved in developmental pathway signaling and growth *(ptgdsb*.*1, avil, mdkb, mdka*), glycogen synthesis (*gyg1b*), innate immune response (*hmgb2a*), vesicle trafficking (*grtp1a*) and markers of goblet cell function (*agr1, muc5*.*1*) were increased. The increased expression of genes involved in the Notch pathway is consistent with our previous finding that the microbiota promotes goblet cell fates through regulation of Notch signaling (Bates *et al*., 2006; Troll *et al*., 2018).

### The microbiota stimulates functional maturation and activation of the immune and neural cells

#### Immune Cells

Immune cells are dedicated to surveilling and responding to microbial cues and previous studies in gnotobiotic zebrafish have illustrated the responses of these cells to microbiota (Bates *et al*., 2006; Kanther *et al*., 2014; Rolig *et al*., 2015; Murdoch and Rawls, 2019; Wiles and Guillemin, 2020). Neutrophils, one of the major immune cell populations in the larval zebrafish, marked by *mpo* segregated to clusters 72. Macrophages, marked by *mpeg1*.*1*, were found in cluster 36 (Fig. 4A&B). We additionally annotated cluster 35 as immune cell progenitors as it showed an enrichment of several genes involved in larval hematopoietic development including *ikzf1* and genes involved in cell cycle regulation and proliferation, such as the proliferation marker *pcna*. Cluster 35 also contains markers of lymphocytes, which start to develop during larval life, including *rag1* and *zap70* (Iwanami *et al*., 2020). In the neutrophil cluster 72, genes enriched in the presence of microbiota encoded characteristic immune cell functions including threat sensing, chemotaxis and cell shape changes, and cellular processing (Fig. 4C). These findings were similar for CVZ compared to GF *mpeg1*.*1*+ macrophages and were cross validated with the Zebrafish Atlas immune cells (Fig. 4C&D). Among the CVZ enriched genes within the progenitor immune cell population of cluster 35 were genes involved in DNA recombination and break repair (S.Clu35_GOsorted_CVZup.xlsx), consistent with the microbiota stimulating lymphocyte maturation.

**Figure 4.**
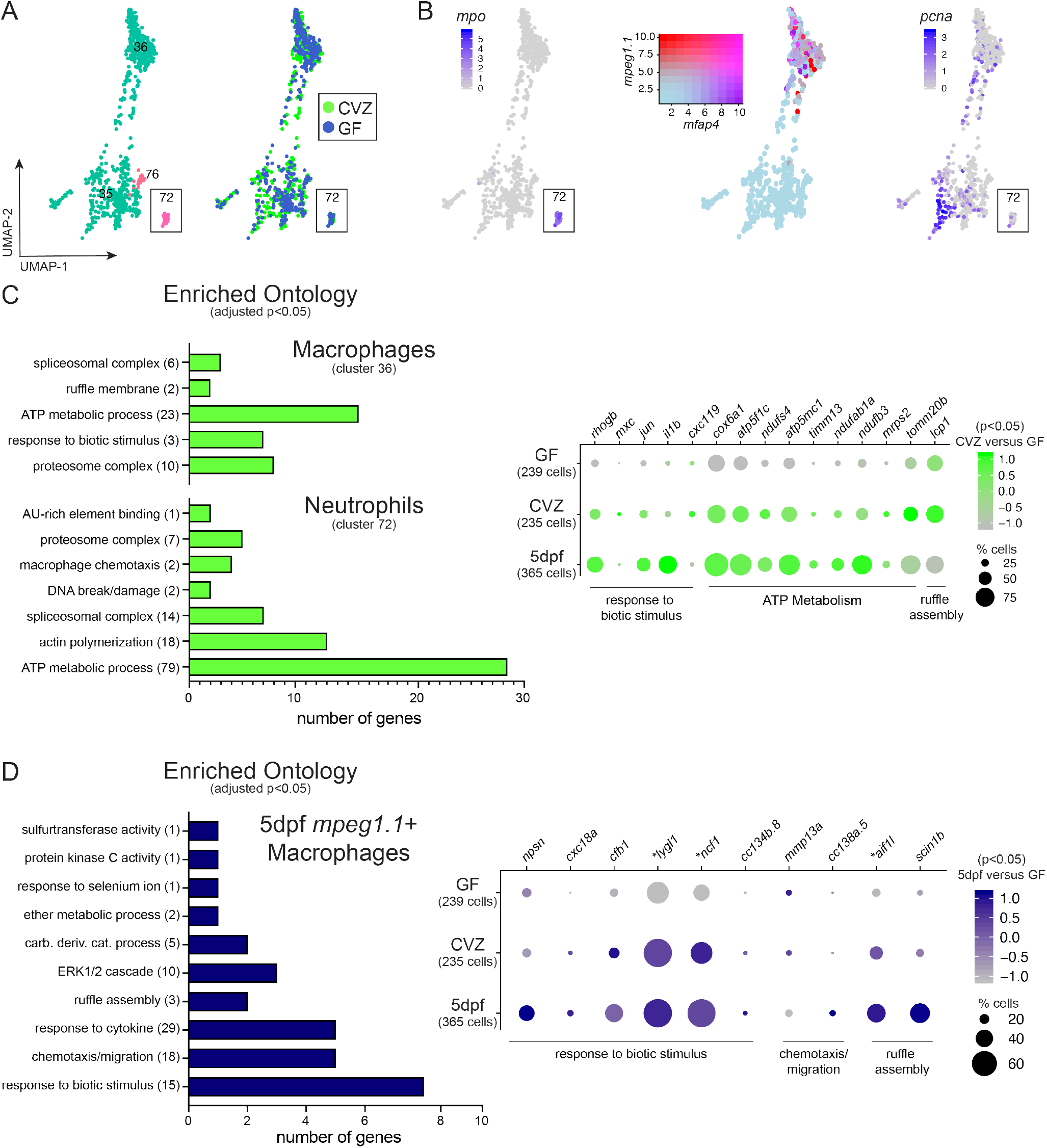
Immune cells exhibit characteristic transcriptional responses to the microbiota. A) Clusters 35, 36, 72 and 76 include several immune cell types that differentially express B) neutrophil biomarker *mpo* and varying degrees of co-expression of macrophage biomarkers *mpeg1*.*1* and *mfap4*. Expression of *pcna* within cluster 35 indicates a population of immune progenitors. C) GO analysis plot and dotplot represent genes significantly enriched within CVZ versus GF macrophages and neutrophils, showing similar trends in gene expression when compared to conventional 5dpf larvae of the Zebrafish Atlas. D) GO analysis plot and dotplot represent genes significantly enriched genes within CV 5dpf Zebrafish Atlas macrophages and neutrophils, showing similar trends in gene expression when compared to 6dpf CVZ cells. Genes designated with * were statistically enriched within CVZ versus GF cells.

#### Enteric Neurons

Mounting evidence indicates that both the peripheral (PNS) and central (CNS) nervous system branches sense and respond to the microbiota (Hsiao *et al*., 2013; Sampson and Mazmanian, 2015; Arentsen *et al*., 2016; Bruckner *et al*., 2020; Jameson *et al*., 2020). Our dataset contained a large number of neurons, marked by expression of the panneuronal, postmitotic marker *elavl4*. Here we focus on intestine-resident neurons of the enteric nervous system (ENS), a major component of the PNS that regulates diverse functions within the intestine including secretion, motility and homeostasis (Spencer and Hu, 2020).

Based on expression of *phox2bb* and *phox2a*, enteric neurons segregated to cluster 33 (Fig. 5A). Cluster 33 is composed of multiple types of PNS neurons as only 52% and 60% of the cells in the cluster express *phox2bb* and *phox2a*, respectively; other top expressed genes in the cluster were identified as biomarkers of cranial ganglia (S.Table1). The co-clustering of enteric neurons and cranial ganglia is consistent with both cell types being derived from neural crest. Re-analyzing cluster 33 alone resulted in 7 subclusters, with high co-expression of *phox2bb* and *phox2a* in subclusters 4, 5, and 6 (Fig. 4B). These subclusters showed expression of transcription factor *hoxb5a*, providing additional evidence they are composed of enteric neurons (Shepherd and Eisen, 2011) (Fig. 4C). Transcriptional differences between these three enteric neuron clusters reflected temporal progression of their maturation (Taylor *et al*., 2016). Subcluster 6 is characterized by expression of early ENS fate determinants (*hoxb5a, hand2*, and *sox10*) and limited neural receptor and enzyme expression, suggesting this population contains enteric neuron progenitors. In subcluster 4, the combinatorial expression of *elavl4* and *ret*, with little expression of *sox10*, suggests a differentiating population (Taylor *et al*., 2016). Subcluster 4 shares some transcription factor expression with subcluster 6, but increased expression of neural receptors (*ngfra, chrna5, chrna3*) and enzymes (*nos1*) is consistent with increasing functionality. Subcluster 5 resembles fully mature enteric neurons lacking expression of *ret* and *sox10* (Taylor et al. 2016) and enriched in expression of neural receptors (*htr3a, ngfra, chrna5*) and neural markers (*calb2a, elavl4*).

**Figure 5.**
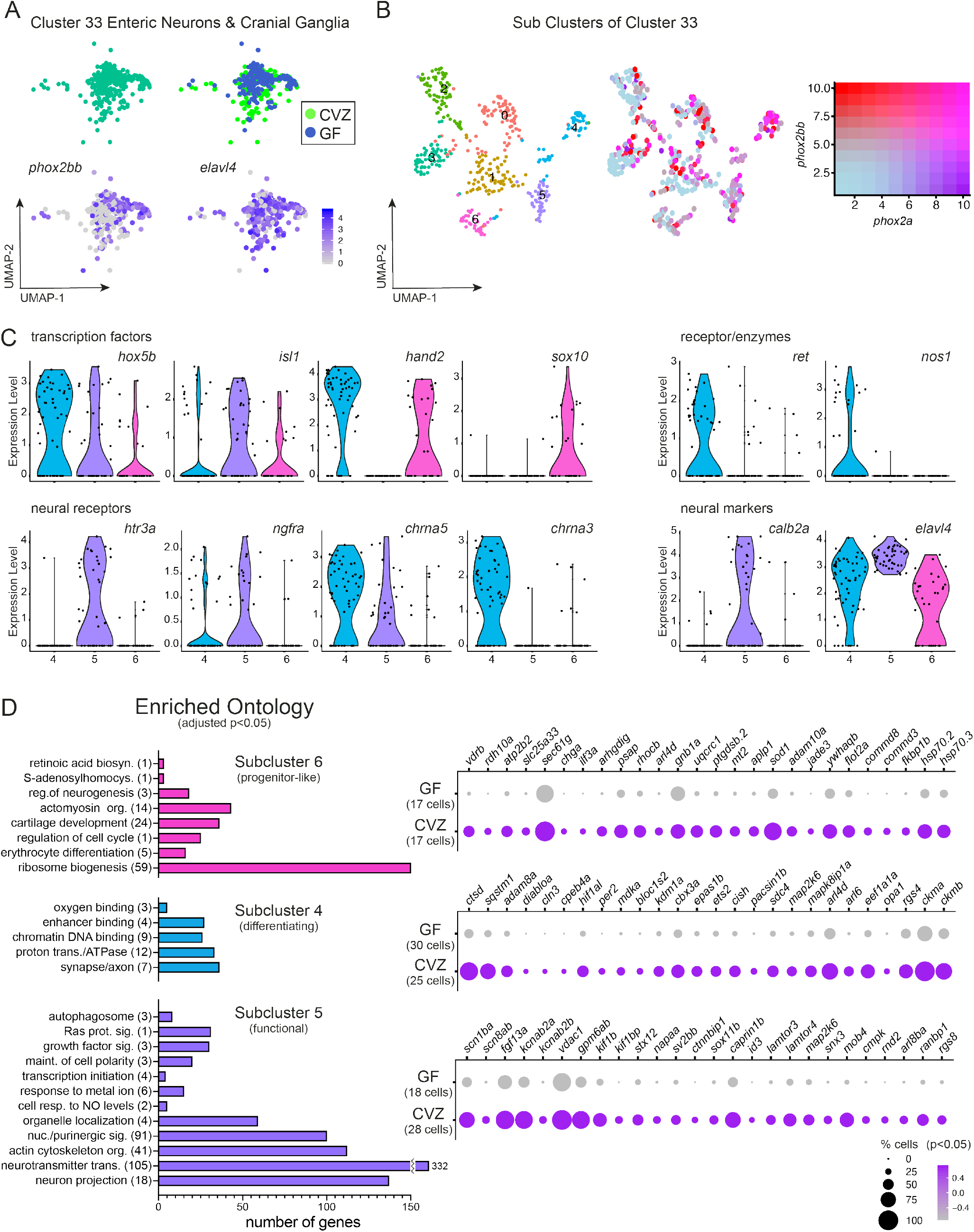
Enteric neurons are transcriptionally heterogenous in response to the microbiota. A) Cluster 33 includes enteric neurons and cranial ganglia showing expression of *phox2bb* and *elavl4*. B) uMAP plots show transcriptional heterogeneity of cluster 33 with a range in co-expression of *phox2bb* and *phox2a*. C) Violin plots compare expression of enteric neuron transcription factors, neural receptors, and neural markers within subclusters 4, 5, and 6. D) GO analysis plots represent genes enriched within subcluster 4, 5 or 6 relative to the other subclusters. The corresponding dotplots represents differential gene expression in CVZ versus GF cells for each subcluster.

To further explore identities of these enteric neuron subpopulations, we performed GO analysis on genes significantly expressed within each subcluster followed by comparing transcriptional differences between CVZ and GF cells for each population (Fig. 5D). Within the progenitor-like population of subcluster 6, several genes involved in ribosome biogenesis, regulation of cell cycle and neurogenesis were enriched versus the other subclusters. This cell population expressed several genes involved in defense response to bacteria and viruses, metal ion binding activity and cell-cell adhesion were enriched within CVZ cells versus GF. Within the differentiating population of subcluster 4, GO analysis revealed an enrichment of genes associated with chromatin DNA binding and the synapse or axon. CVZ cells of subcluster 4 also showed enrichment of genes involved in defense response to microbes, as well as genes involved in cell maintenance and differentiation. GO analysis of genes enriched within subcluster 5 is consistent with mature neurons including: maintenance of cell polarity, purinergic signaling, neurotransmitter transport and axon projection. The presence of the microbiota induced the enrichment of many genes in CVZ cells including those involved in voltage-gated ion channel activity and neurotransmission. Our analysis of enteric neurons corroborates their transcriptional heterogeneity (Taylor *et al*., 2016) and illustrates how cells of different maturation states respond differently to the microbiota.

#### Central Neurons

Our analyses of CNS neurons also showed transcriptional responses to the microbiota that differed between cells in different maturity states. We identified a neuronal population likely to be composed of progenitors within cluster 21 by expression of the nervous system development gene *ptn* and co-expression of *fapb7a* and *her4*.*2* (S.Fig. 8A&B) (Farnsworth, Saunders and Miller, 2019). Similar to progenitor-like enteric neurons, several genes involved in regulation of neurogenesis were enriched within cluster 21 CVZ neurons, as were genes involved in immune function and development (S.Fig. 8C&D). These findings are consistent with mounting evidence implicating connections between immune response and neurodevelopment (Pronovost and Hsiao, 2019). We additionally observed that mature serotonergic and/or dopaminergic neurons in cluster 69 responded to the microbiota by upregulating genes involved in synaptic machinery for dopamine transmission (S.Fig. 8E&F). CVZ neurons of cluster 69 also exhibited significant enrichment of genes involved in MAP kinase activity, negative regulation of proliferation, and calcium binding activity, suggesting that the microbiota plays a role in promotes characteristic function of mature neurons (S.Fig. 8G). Together these data suggest the microbiota plays specific and distinct roles within neuronal cell types, depending on their developmental state, and promotes neural functions at all developmental stages, including neurogenesis, neurodifferentiation, and neurotransmission.

#### Host tissues exhibit global patterns of microbiota responsiveness

Our GO analyses across each of the 78 clusters revealed two striking patterns. First, ATP metabolism genes such as *atpf51b, COX5B, vdac2*, which were widely expressed across all cells, were consistently expressed at higher levels in cells from CVZ animals. Second, whereas lens-associated *crystallin* (*cry*) genes were almost entirely restricted to the lens cell clusters 39 and 55 in CVZ animals, there was widespread expression of multiple *cry* genes across virtually all GF clusters, predominantly of the beta and gamma type (Fig. 6A-C, SFig. 9). We explored each of these transcriptional signatures further.

**Figure 6.**
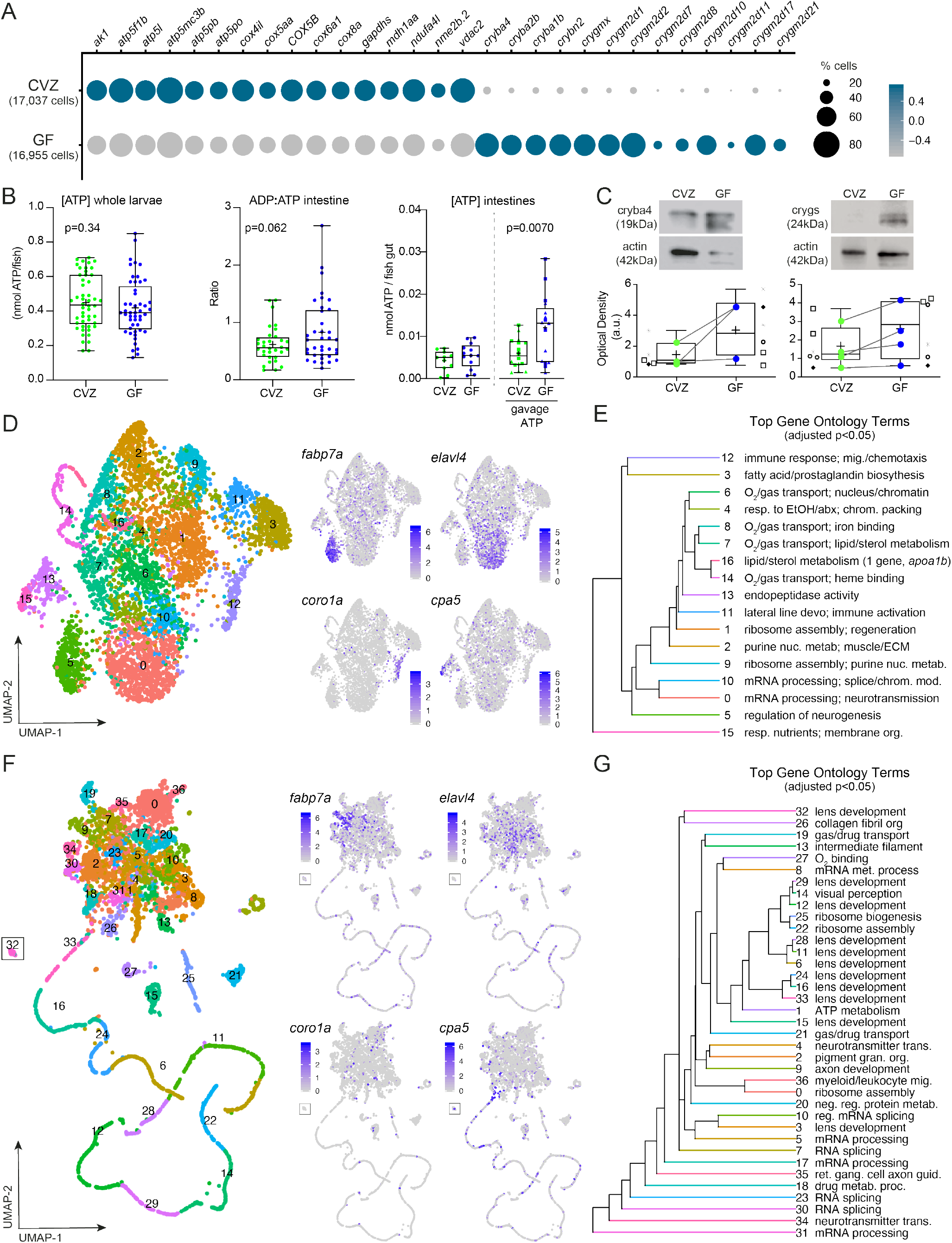
The microbiota induces global patterns of host gene expression. A) Dotplot illustrates global trends of gene expression across all CVZ and GF cells from whole larvae. B) Boxplots show the concentration of ATP from whole larvae, the ratio of ADP:ATP of dissected larval intestines, and the concentration of ATP in dissected CVZ and GF larval intestines 20-40 minutes following gavage with 1 or 10mM ATP. Each dot in the measurement of ADP:ATP represents the average to two intestines. C) Boxplots show the average relative optical density of beta and gamma crystallin protein expression in CVZ and GF larvae. Each symbol represents an individual flask of larvae processed together and matching symbols denote that flasks were from the same gnotobiotic derivation. Colored dots represent average expression between flasks in the same derivation with a line connecting the corresponding derivations between experimental groups. D) uMAP plots represent CVZ cell clustering based on expression of genes involved in ATP metabolism. Feature plots illustrate cell types maintaining their original cell type identity. E) Hierarchal tree demonstrates the proximity of uMAP plot cells clustered by expression of ATP metabolism genes and the top GO terms based on enriched genes within a given cluster. F) uMAP plots represent corresponding GF cells from the original clusters used in D’ clustering based on expression of crystallin genes. Feature plots show that clusters lose their original cell type identity. G) Hierarchal tree demonstrates proximity of clusters from the uMAP plot of cells clustered by expression of *crystallin* genes and the top GO terms based on enriched genes within a given cluster. Data from *in vivo* experiments, and for the remaining figure, are shown as boxplots with the middle line representing the median, the top and bottom of the box representing the upper and lower quartile, the whiskers representing the min and max values, and the ‘+’ symbol representing the mean. Each dot symbol represents data from a single 6dpf zebrafish, unless otherwise specified. p-values displayed on boxplots are from Students’ Ttest, where p<0.10 is considered trending and p<0.05 is considered significant.

#### The microbiota elicits cell type specific ATP metabolism gene expression patterns

ATP is an ancient molecule whose production spurred one of the earliest examples of host-microbe interactions with the endosymbiosis event that produced mitochondria (Margulis, 1976). ATP production has been shown to be modulated by the microbiota in many model organisms. In GF fruit flies, whole body ATP levels and mitochondrial respiration are significantly lower than in conventional counterparts (Gnainsky *et al*., 2021). In mice, ATP levels and mitochondrial respiration are reduced in colonic tissue (Donohoe *et al*., 2011) and lower in feces from GF animals (Atarashi *et al*., 2008). Enriched expression of respiratory electron transport chain genes involved in ATP metabolism was also observed in dissected 6dpf digestive systems of CVZ versus GF zebrafish larvae (Willms et al. 2022). We measured the concentration of ATP in whole zebrafish larvae and observed extensive inter-individual variation in levels of this rapidly metabolized molecule with no significant difference between GF and CVZ groups (Fig. 6B). We explored ATP turnover in dissected larval intestines and found that ADP:ATP ratios varied extensively across individuals, with a trend toward lower ratios in CVZ larvae. We tested the rate of ATP metabolism in the presence and absence of microbiota by microgavaging exogenous hydrolysable ATP into the proximal intestinal lumen, and 20-40 minutes later dissecting out the intestines and measuring ATP levels. ATP levels were significantly lower in CVZ samples, indicating that ATP was consumed faster within CVZ intestines. These measurements are consistent with the transcriptional signature of upregulation of enterocyte cellular activities.

ATP production for energy currency or cell-signaling can vary extensively across diverse cell types. For example, neurons use massive amounts of ATP for both neural activity (MacVicar, Wicki-Stordeur, and Bernier 2017) and as a signal co-released with various neurotransmitters (Huang, Otrokocsi, and Sperlágh 2019). Neutrophil production of ATP sustains their immune cell activation and also initiates purinergic signaling for chemotaxis (Bao *et al*., 2014). Murine colonic macrophages have been shown through single cell transcriptomic analysis to upregulate oxidative phosphorylation genes in response to the microbiota (Kang *et al*., 2019). One third of the clusters in our dataset, representing diverse cell types, exhibited enriched gene ontologies pertaining to nucleotide and/or ATP metabolism in the CVZ versus GF cells (S.Fig. 10). To better understand how ATP metabolism genes are deployed in response to microbiota, we bioinformatically isolated CVZ cells from the clusters with enriched ATP metabolism genes and re-clustered them using a panel of 228 ATP metabolism-associated genes (Fig. 6D). This analysis revealed 17 new subclusters that largely maintained their original cell-type identity, including neural progenitors (84% of original cluster within subcluster 5, *fabp7a*), neurons (73% of original cluster within subcluster 0, *elavl4*) and immune cells (88% of original cluster neutrophils, within subcluster 12, *coro1a*) (S.Table4). Some original populations were split into different subclusters, such as acinar cells of original cluster 32 exocrine pancreas with 35% within subcluster 15, 19% within subcluster 13 and 14% within subcluster 0, suggesting heterogeneity of exocrine pancreas ATP metabolism gene utilization. GO analysis illustrated a diversity of biological processes among the enriched genes within each subcluster, consistent with cell type specific functions (e.g. upregulation of chemotaxis in neutrophils). This analysis demonstrates that the microbiota stimulates cell type specific ATP metabolism gene expression.

#### The microbiota suppresses cry gene expression across tissues

Crystallins are a heterogenous group of extremely stable proteins that make up the vertebrate eye lens. In most vertebrates, including zebrafish, rodents, and primates, the lens is comprised of alpha crystallin small heat shock proteins and the beta and gamma crystallin superfamily members. The zebrafish genome contains 3 alpha *cry* genes, 13 beta *cry* genes and 40 gamma *cry* genes (Farnsworth et al., 2021). Related *cry* genes were present in early vertebrates prior to lens evolution, implying that they have additional functions (Zigler and Sinha, 2015). In both rodents and zebrafish, alpha and beta crystallin are expressed and function in other tissues beside the lens, including the retina, brain, heart, and testes (Piri, Kwong and Caprioli, 2013; Zigler and Sinha, 2015; Sprague-Piercy *et al*., 2021). At the protein level, we confirmed expression of Cryba4 is not restricted to the head or lens of GF larvae but is also detected within the body (S.Fig. 11A). Within whole larvae across several GF derivations, the levels of Cryba4 and Crygs protein were consistently increased in samples of pooled GF larvae (Fig. 6C). We note that we did not observe ectopic *cry* transcripts in dissected digestive system cells from GF larvae, nor did Willms et al. 2022 report an enrichment of *cry* transcript expression in their GF cells from dissected digestive tracts. We suspect that the additional processing of these samples, including incubation of the tissue in enriched media, attenuate *cry* gene expression.

Alpha crystallins function as chaperones that prevent aggregation of client proteins damaged by heat or oxidative stress (Sprague-Piercy *et al*., 2021). Alpha crystallins have antiapoptotic and neuroprotective functions in the retina and other neuronal tissues (Zhu and Reiser, 2018; Phadte, Sluzala and Fort, 2021). In the retinal epithelium, both alpha and beta crystallins stabilize the vacuolar-ATPase, thereby protecting against lysosomal dysfunction (Valapala *et al*., 2016; Cui *et al*., 2020). In several mouse models of maternal immune activation, there is a transient upregulation of representative members of all *Cry* gene families within the embryonic brain, consistent with neuroprotection (Garbett *et al*., 2012). In zebrafish, Cryab confers protection against cardiac stress induced by crowding or cortisol (Mishra *et al*., 2018).

We explored whether *cry* genes were co-expressed with other chaperone genes in GF cells. Although we saw strong co-expression of *cry* genes with each other, they were not co-expressed with heat shock genes *hsp70*.*2* and *hsp70*.*3* (S.Fig. 10B). We also did not observe any relationship between the expression of crystallin genes and ATP metabolism genes (S.Fig. 10B). We next explored how *cry* genes are regulated in the absence of microbiota by re-clustering GF cells based solely on the 87 *cry* genes found within the sequencing data (Fig. 6F). In contrast to clustering with ATP metabolism genes, this analysis revealed clusters that largely lost their original cell-type identity; for example, neural progenitors (*fabp7a*), postmitotic neurons (*elavl4*), and immune cells (*coro1a*) no longer co-segregated within a specific sub-cluster (S.Table5), showing that *cry* genes are expressed across many cell types in the absence of the microbiota.

Among the GO analysis of GF cells clustered by *cry* genes, reoccurring themes were related to cellular architecture (e.g. collagen fibril organization and intermediate filaments) and vesicle trafficking (e.g. neurotransmitter transmission and gas/drug transport) (Fig. 6G). Comparing differentially expressed genes across all CV and GF cells showed that in addition to *cry* genes, several digestive enzymes were enriched within 10-40% of GF cells (S.Fig. 12A). These digestive enzymes include *prss59*.*1* and *ela2*, which were enriched within 6dpf GF dissected digestive systems (Willms *et al*., 2022). We also observed significant co-enrichment of several *keratin* genes and *cry* genes in GF cells (S.Fig. 12A-C), which was more pronounced in GF *elavl4*+ cells (predominantly postmitotic neurons) (S.Fig. 12D). Keratins are intermediate filaments that confer structural integrity to postmitotic cells. Intermediate filaments are critical for neural development but their myriad functions have been underappreciated (Bott and Winckler 2020), such as stabilizing microtubules and inhibiting synaptic vesicle trafficking during synaptic rewiring in postmitotic neurons (Kurup *et al*., 2018). We hypothesize that widespread expression of *cry* genes in GF cells counteracts cellular stresses induced by the artificial GF state. Beyond their roles in preventing lens protein aggregation and cataracts, Cry proteins chaperone cytoskeletal components and maintain cytoplasmic organization in diverse postmitotic cells (Slingsby and Wistow, 2014). In a meta analysis of zebrafish proteins upregulated in response to various biological stressors, Cry proteins are among the top 25 protein families (Groh and J-F Suter, 2015). More broadly, small heat shock proteins similar to alpha crystallins are abundant in cells experiencing low metabolic states, such as in nematode dauers (Fu *et al*., 2021) and brine shrimp and insects in diapause (King and Macrae, 2015; Khodajou-Masouleh *et al*., 2021). Our global transcriptional analysis of GF cells suggests they are less proliferative and less metabolically active than CV counterparts, potentially necessitating cellular mechanisms to maintain cytoplasmic organization, prevent protein aggregation, and stabilize stalled, energetically costly processes, such as vesicular trafficking.

#### Exocrine pancreas responses illustrate how the microbiota promotes tissue development and function

Of all the cell clusters, the exocrine pancreas acinar cell cluster 32, marked by high expression of digestive enzymes including *amy2a* and *cpa5*, had the largest number of microbiota-induced genes (Fig. 1D). The exocrine pancreas produces digestive enzymes that are packaged into vesicle-granules and delivered into the intestinal lumen by secretion. GO analysis indicated that the microbiota stimulates a diversity of biological processes in exocrine pancreas acinar cells including secretory vesicle formation, transporter activity, and regulation of catabolism, consistent with several studies showing associations between exocrine pancreas function and microbiota composition (Nishiyama *et al*., 2018; Adolph *et al*., 2019; Frost *et al*., 2019; Pietzner *et al*., 2021). Pancreatic acinar cells are not only responsible for organ function but also for organ development (Biemar *et al*., 2001; Field *et al*., 2003; Wan *et al*., 2006). Other genes upregulated within CVZ acini included genes involved in DNA-binding transcription factor activity (*ebf3a, bcl11ba, sox4a*) as well as growth factor activity and organ development (*mdkb, ppdpfb, fbxl3a, gng2, ptmaa, akt1s1, serinc1*). Specifically, enriched expression of *ppdpfb* and *mdkb* within CVZ cells suggests that the microbiota promote pancreas development.

In contrast to the increased expression of genes involved in acinar cell differentiation and function in CVZ cells, we noted a subtle but consistently elevated level of digestive enzyme transcripts in GF acinar cells (Fig. 7B). Expression of these digestive enzyme genes was uniformly high in GF acinar cells, whereas CVZ acini shows a bi-modal distribution, consistent with different states of differentiation and maturation. To validate transcriptional increases of digestive enzymes within GF exocrine pancreas, we derived Tg(*ptf1a*:GFP) zebrafish CVZ and GF to mark the exocrine pancreas and stained larvae with an antibody against Amylase to characterize expression of this digestive enzyme (Fig. 6C&D). GF larvae had higher intensity of Amylase labeling within the pancreas (Students’ Ttest, p=0.0019) but the average pancreas volume was larger within CVZ larvae (Students’ Ttest, p=0.0196). These analyses reveal how microbiota stimulate both tissue growth and function, promoting developmental plasticity in response to the microbial environment. The transcriptional plasticity of pancreatic cells has also been demonstrated at single cell resolution within regenerating islets of the zebrafish endocrine pancreas (Singh *et al*., 2022). In the artificial GF state, the exocrine pancreas appears to be stunted and composed of fully differentiated acinar cells that, based on their transcriptional patterns, are poised in a functionally inactive state, a situation that may require upregulation of cytoprotective Cry proteins to maintain.

**Figure 7.**
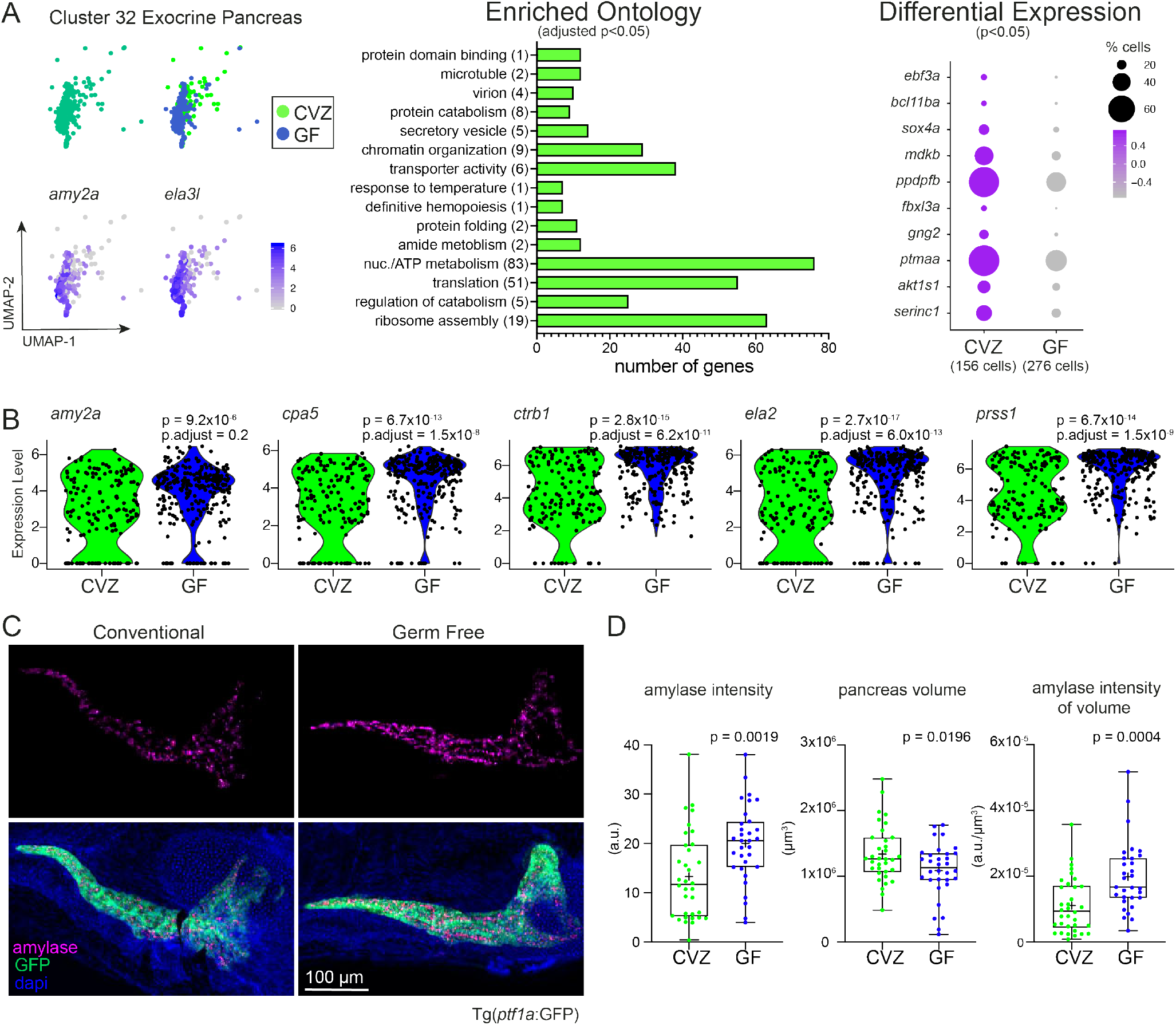
The microbiota promotes tissue development and function within the exocrine pancreas. A) Cluster 32 is composed of acinar cells from the exocrine pancreas showing high expression of digestive enzymes *amy2a* and *ela3l*. GO analysis plot and genes included in the dotplot are based on genes significantly enriched within CVZ cells of cluster 32. B) Violin plots illustrate the difference in distribution of pancreatic digestive enzyme gene expression between CVZ and GF cells from cluster 32. C) Images display amylase protein expression within the exocrine pancreas of CVZ and GF larvae. Images are z-projections taken by tile-scan with a confocal microscope. D) Boxplots illustrate the average optical intensity of amylase expression, the total volume of space occupied by the pancreas, and the intensity of amylase standardized to the pancreas volume. All images were taken with the same optical settings and larvae from each experimental group were imaged on the same day in the same session. Size bar of 100 µm corresponds to all micrographs.

## Conclusion

Our analyses reveal widespread and diverse host responses to the microbiota that are not limited to host cells in direct contact with resident microbes. We found that exposure to microbes simultaneously stimulates cell type specific programs of microbial responses and metabolic activity, highlighting the close links between immunity and metabolism (Hotamisligil, 2017). The presence of microbes stimulates cellular programs of proliferation and regeneration in diverse progenitor-like cell types, demonstrating the importance of microbial cues in organismal development. In addition, microbes stimulate the cellular activities of fully differentiated cells from protein-ingesting enterocytes to dopamine-secreting neurons. In contrast, organisms raised in the absence of microbes express unusual repertoires of Cry proteins, suggesting a requirement to stabilize the cytoplasmic organization of fully differentiated but metabolically inactive cells. Collectively our profiling of the cellular transcriptomes of CVZ and GF developing animals reveals myriad roles of resident microbes in stimulating tissue development and cellular functions, demonstrating how microbiota composition and assembly can shape organismal development in a manner that is plastic to the environment the adult organism will inhabit. Our study illustrates the utility of single cell transcriptomics for exploring intricate interactions between vertebrate hosts and their resident microbes and provides a valuable resource for generating new hypotheses about the impacts of the microbiota on animal development and physiology.

## Supporting information

Lists of DEGs and GO per Cluster

## Acknowledgements

We thank C. Robinson, E. Wall, and T. Wiles for assistance with GF derivation, J. Hill for assistance with dissections, M. Weitzman, D. Turnbull and the UO Genomics Core for assistance with transcriptomics, P. Batzel, H. Healey, A. Kershner, G. Perez, N. Tran, and T. Wiles for assistance with analysis, and R. Sockol and the University of Oregon Zebrafish Facility for zebrafish husbandry. Research reported in this publication was supported by the NIH under award numbers 1P01GM125576 (to K.G.) and 5T32HD007348 (to M.S.M.).

## Author Contributions

Conceptualization, M.S.M., K.G., Software, M.S.M., G.K., D.C., Validation, M.S.M., Analysis, M.S.M., G.K., D.C., M.K.H., E.M., Investigation, M.S.M., Resources, K.G., Data Curation, M.S.M., G.K., D.C., Writing-Original Draft, M.S.M., K.G., Writing-Review & Editing, M.S.M., K.G., J.S.E., Visualization, M.S.M., Supervision, K.G.

## Figure Legends

**Supplemental Figure 1. Determination of principle components for downstream analyses**. A) Jackstraw plot illustrates the p-value for each principle component (PC) and B) Elbow plot displays how much variation is attributed to each subsequent PC. C) Heatmap of genes significantly contributing to variation within PC1 are those expressed within the liver and integument.

**Supplemental Figure 2. Inclusion of additional PCs increases the number of clusters**. Heatmap, resulting uMAP, and expression of intestinal biomarker *vil1* are displayed after inclusion of the first A) 30, B) 60, C) 90, and D) 122 PCs.

**Supplemental Figure 3. Transcriptomic profile of 6dpf CVZ cells are similar to 5dpf larvae of the Zebrafish Atlas**. A) Integration of the Zebrafish Atlas 1, 2, 5dpf whole organism single cell RNAseq data (Farnsworth et al., 2019) with CV 6dpf data illustrates that 6dpf cells align more closely with the 5dpf cells than earlier timepoints, with the exception of neural cells (*elavl4+*) which are B) largely absent within the 5dpf data sets, possibly due to differences in cell dissociation protocols.

**Supplemental Figure 4. GF zebrafish lack representation of an epithelia cell type**. A) uMAP plots display a transcriptionally distinct type of epithelium is overrepresented within CVZ versus GF cells despite different permutations of PC inclusion. B) Cluster 45 has high expression of *krt17* and *cyt1* similar to cluster 19 epithelia. C) GO analysis plot is based on genes enriched within cluster 45 versus all other clusters and D) dotplot illustrates that the top markers expressed within cluster 45 are broadly expressed across different clusters that are categorized as epithelia. The * designates clusters that contain cells from the intestinal epithelium.

**Supplemental Figure 5. Dissected intestines of larval zebrafish contain a diversity of digestive system cell types**. A) Digestive systems of larval zebrafish were dissected, cells were dissociated and subjected to single cell RNA sequencing. B) uMAPs show diversity and enrichment of enterocytes and associated digestive system cells.

**Supplemental Figure 6. Comparison of differentially expressed genes in gnotobiotic microarray versus single cell RNA-sequencing of digestive systems**. Common genes between dissected gnotobiotic intestines of 6dpf zebrafish larvae from Rawls et al., 2004 microarray analysis and cell types from single cell dissociations likely included in dissected digestive systems. Congruent genes highlighted in green indicate a similar enrichment within the respective CVZ treatments whereas genes highlighted in red indicate the opposite trend, increased expression within GF single cells. Comprehensive gene expression comparisons between gnotobiotic groups across Rawls et al., 2004 and Willms et al., 2022 dissected digestive systems scRNAseq data is included in S.Table2.

**Supplemental Figure 7. Enterocytes transcriptionally segregate by proximal to distal localization along the intestine**. A) Cluster 11 and 64 are composed of intestinal enterocytes with enriched expression of *vi11, muc13b, fabp6* and *anxa4*. B) uMAP plots show transcriptional heterogeneity of intestinal enterocytes from cluster 11 and 64 which C) segregate by expression of proximal to distal intestinal epithelial biomarkers. D) uMAP plots shows minimal co-expression of secretory cell biomarkers within cluster 11 and 64. E) subcluster 7 has the most consistent expression of secretory cell biomarkers; GO analysis plot is based on genes enriched within subcluster 7 versus other subclusters (pink bars) as well as genes enriched within CVZ cells of subcluster 7 versus GF (green bars).

**Supplemental Figure 8. CNS neurons respond to the microbiota**. A) Cluster 21 is composed of neural progenitors showing enriched expression of progenitor marker *ptn* and B) co-expression of *fapb7a* and *her4*.*2*. C) GO analysis plot and D) dotplot based on enrichment of genes within CVZ versus GF cells of cluster 21. E) Cluster 69 is populated with serotonergic and dopaminergic neurons displaying expression of *ddc, slc18a2, tph1a*, and *th2*. F) GO analysis plot and G) dotplot based on the enrichment of genes within CVZ versus GF cells of cluster 69.

**Supplemental Figure 9. Crystallin genes are enriched within GF cells**. A) Violin plot shows expression of *crygn2* within lens fiber clusters 39 and 55 with enriched expression in GF cells across all clusters. B) uMAP plots compare global expression of *crystallin* in CVZ and GF cells.

**Supplemental Figure 10. The microbiota promotes expression of genes involved in ATP and nucleotide metabolism in diverse cell types**. uMAP highlights clusters composed of diverse cell types that show enrichment of gene expression involved in ATP metabolism within CVZ versus GF cells.

**Supplemental Figure 11. Crystallin expression occurs outside of the lens in GF larvae and is uncorrelated with heat-shock and ATP metabolism genes**. A) Western blot showing protein expression of Cryba4 within heads versus corresponding bodies of GF larvae. Images display expression of Cryba4 in different tissue structures of the larval tail. B) Heatmap plot showing Pearson Correlational coefficients across all CVZ and GF cells in the experiment with respect to *cryba4* and *crygn* to heat-shock and ATP metabolism genes. Heatmap also displays Pearson Correlational coefficients in analyses that only include cells positive for both transcripts.

**Supplemental Figure 12. Co-expression of crystallin and select keratins are enriched within GF cells**. A) Dotplot features transcripts significantly enriched within GF cells accounting for all cells in whole larvae. B) uMAP plots from corresponding GF cells from the original clusters used in Fig.6D clustered based on the expression of crystallin genes show that specific keratins (*cyt* and *krt5*) are broadly expressed across GF cells compared to epidermal specific keratin (*krt17*) that maintains its original cell type identity. C) Corresponding uMAP plots show co-expression of *cyt1* and *krt5* with *cryba4* across GF cells. D) uMAP plots show CVZ and GF *elavl4+* cells separate based on their experimental group when clustered by the top variable gene expression and that co-expression of *cyt1* and *krt5* with *cryba4* is enriched within GF *elavl4*+ cells. E) uMAP plots show that *cyt1* is co-expressed with *cryba4* across diverse cell types within whole larvae. F) Scatterplots demonstrate the correlation of *cryba4* and *ela2* across cells in the dataset that express both transcripts and without cells from the exocrine pancreas cluster 32 and lens clusters 39 and 55).

## Methods

### Gnotobiotic Zebrafish

All protocols used for zebrafish experiments were approved by the University of Oregon Institutional Care and Use Committee. Adult zebrafish were maintained using standard husbandry procedures (Westerfield, 2007) by the University of Oregon Zebrafish Facility. Zebrafish embryos were derived germ free (GF) as previously described (Melancon *et al*., 2017). Conventionalized (CVZ) zebrafish were inoculated with parental tank water post GF derivation. Animals used for the single cell analysis of whole larvae were *Tg*(*nkx2*.*2a*:EGFP) (Pauls *et al*., 2007) and multiple clutches were collected from natural mating of adults from the same tank. The *Tg*(−1.0*ins*:EGFP) (DiIorio *et al*., 2002) line of zebrafish were used for dissections of the digestive system prior to single cell dissociation. Animals used for western immunoblots were wild-type (ABCxTu strain). The Tg(*ptf1a*:EGFP) (Godinho *et al*., 2005; Dong *et al*., 2008) line of zebrafish were used for all analyses with the exocrine pancreas. Unless specified, all zebrafish used were 6 days post fertilization (dpf).

### Whole Larval Zebrafish Dissociation

Larval zebrafish were anesthetized with Tricane (Western Chemical, Inc., Ferndale WA). 15 healthy zebrafish with inflated swim bladders were selected for dissociation and put into a sterile 1.5 ml Eppendorf tube. Stock solutions of 10mg/mL proteinase K (Millipore Sigma, St. Louis, MO) and 10mg/mL collagenase P (Millipore Sigma, St. Louis, MO) were prepared by dissolving in 1X HBSS (Thermo Fisher Scientific, Waltham, MA). Embryo media was pipetted out from each tube and 37C-warmed 1.3 ml of dissociation solution (0.12 mg/mL proteinase K and 1 mg/mL collagenase P in 1x TrypLE (Thermo Fisher Scientific, Waltham, MA) was added to each tube. Each tube was quickly added to a heat block at 37C, with a timer starter. To dissociate the whole 15 zebrafish per tube into individual cells, each tube was mixed by pipetting 25x every 2 minutes until ~12 minutes is on the timer. To stop the dissociation reaction, 200 ul of 4C ‘stop solution’ (5% calf serum, 1 mM CaCl2, and PBS) (Farnsworth, Saunders and Miller, 2019) was added to each tube, mixed by pipetting 5x, and immediately put into a pre-chilled 4C table-top centrifuge. Dissociated cells were gently spun down for 3 minutes at 350g and the supernatant removed. Cell pellets were gently rinsed and resuspended with 1mL of 1% Bovine Serum Albumin (BSA) in HBSS and pelleted again for 3 minutes at 350g. The supernatant was removed and the pellet resuspended in 100 µl of 0.04% BSA/HBSS. Cells were then filtered through a pre-chilled 40 µM strainer into a pre-chilled Eppendorf tube, using a pre-chilled 1mL syringe plunger to gently pestle the cells through. An additional 100 µl of fresh 0.04% BSA/HBSS was used to rinse remnants of the strainer into the tube. A Bio Rad TC20 cell counter was used to measure cell concentration and cell viability with Trypan Blue (Bio Rad, Hercules, CA). Additionally, pilot experiments confirmed single cell dissociation with this protocol by visual inspection using the DIC feature of a LEICA DM6 confocal microscope (Leica Microsystems Inc., Buffalo Grove, IL). All groups of dissociated cells had over 80% viability and were diluted to a final concentration of 3,500 cells/ul in 0.04% BSA/HBSS for cDNA library preparation. The timing from euthanasia to handing off dissociated cells for library preparation was under 30 minutes.

### Larval Zebrafish Dissections & Dissociation

For larval zebrafish dissections, the entire digestive system (intestine, pancreas, liver) was dissected as previously described (Rolig *et al*., 2015). Briefly, zebrafish were derived germ free and at 5dpf were anesthetized in Tricane (Western Chemical, Inc., Ferndale WA), mounted on a slide and their digestive systems sterilely dissected. Dissected tissue was isolated and put into L-15 culture medium (Thermo Fisher Scientific, Waltham, MA) supplemented with 10% fetal bovine serum (Thermo Fisher Scientific, Waltham, MA), penicillin-streptomycin (MilliPore Sigma, St Louis, MO) and gentamycin (VWR, Radnor, PA). Dissected tissue was incubated overnight in culture media at room temperature prior to single cell dissociations.

### Single-Cell cDNA Preparation

The University of Oregon Genomics and Cell Characterization Core Facility https://gc3f.uoregon.edu/ performed the sample preparation by running the samples on a 10X Chromium Single Cell 3’ platform using v2 chemistry. The goal was to target 10,000 cells per group and the resulting cDNA libraries were amplified with 10 cycles of PCR. The cDNA libraries were first sequenced with experimental groups combined onto a single Hi-seq 4000 lane. This resulted in a low coverage depth. We next had each experimental group sequenced onto its own Hi-seq 4000 lane. The output from all lanes was combined, which reached an optimal number of reads for each sample. All samples were prepared and sequenced on the same days.

### Computational Analysis

The sequencing data was aligned to the zebrafish genome (GFCz11_91) using the 10X Cell-ranger pipeline (3.0.2). The Seurat software package for R (3.1.4) was used to subject the data to standard pre-processing workflow prior to integrating CVZ and GF cells together. The data was filtered such that any cells expressing more than 3,000 genes, including more than 50,000 read counts, or more than 20% mitochondrial genes were not included in the final analysis. The expression level of each gene was normalized by total expression via log-transformation with a 10,000 scale factor. We performed a linear dimensional reduction of the data by principle coordinate analysis (PCA) and calculated 150 principle components (PC). Based on Jack Straw methods to determine significance of PCs, the first 122 PCs significantly explain the variance of the data (p<0.0001) but by Elbow Plot most of the variation can be explained between 30 to 60 PCs (S.Fig. 1). To better grasp how adding additional PCs impact our data, we performed multiple clustering analyses comparing the inclusion of 30, 60, 90, and 122 PCs (S.Fig. 2) using all genes in the dataset. The data we are reporting in subsequent figures includes 60 PCs with a resolution of 3.0, which produces 78 clusters where we can conservatively identify distinct cell types and avoid possible technical noise.

Gene expression conserved between the gnotobiotic treatments for all the clusters was performed using the FindConservedMarkers function in Seurat. The lists of genes that are conserved between CV and GF treatments and significantly contribute to the segregation of cells into each cluster is made available in the workbook, S.Table 1. We then isolated the data from specific cell types by sub-setting a cluster(s) or subset cells globally that were positive for a biomarker(s) of interest. Re-analyzing specific cell type populations allowed us to understand the extent of gene expression heterogeneity within a cell population and the FindMarkers function was used to find differentially expressed genes between CV and GF cells in the sub populations by Wilcoxon rank sum tests. These analyses took into account the expression of all present transcripts within the cells prior to clustering except for the analysis of *elavl4+* cells which used the top variable genes per the standard workflow in Seurat. Gene ontology (GO) analysis on the lists of differentially expressed genes (p<0.05) was done using the ClusterProfiler software package for R (3.14.3) and the genome wide annotation for zebrafish org.Dr.eg.db (3.10.0). In the GO analyses, input genes are assigned a GO term and are bioinformatically associated with GO annotations, which provides a statement about the function or biological context of genes based on current biological knowledge http://geneontology.org/. Resulting GO terms were subject to FDR p-value correction to limit potential false-positives. To distill the repetitive GO annotations and terms, we manually triaged through each analysis included in the figures and binned similar annotations that were described by the same sets of genes into broader categories. Within figures, GO analyses are shown as bar graphs displaying the number of cumulative genes that contribute to the GO category. All GO terms, genes that contribute to the terms and which terms were binned into our broad categories is made available as supplemental spreadsheets.

### Western Immunoblot

Larval zebrafish were euthanized by tricane and ~30-40 zebrafish were added to an Eppendorf tube. Leftover embryo media was removed from each tube and replaced with 400-600 µl of 4C from a 10mL stock of cold lysis buffer (RIPA buffer (Boston BioProducts, Ashland, MA) and ½ protease cocktail inhibitor (Thomas Scientific, Swedesboro, NJ)). Samples were sonicated with a microtip at 20% amplitude, 1 second pulse on, 0.3 second off for 30 seconds. This step was repeated until fish were completely dissociated, taking care that a sample would not become warm. Tubes were then put in the freezer for 15 minutes, followed by thawing on ice prior centrifugation at 14,000 rpm for 20 minutes at 4C. The supernatant from each tube was isolated and protein concentration for each sample was quantified using the Pierce BCA Protein Assay Kit (Thermo Scientific, Rockford, IL). 20 µg of each sample was loaded onto a 4-20% Gel (Bio Rad, Hercules, CA) for electrophoresis and transferred to a PVDF membrane (GE Healthcare Amersham, Chicago, IL). For cryba4 (Thermo Fisher Scientific, Waltham, MA) detection, membranes were blocked in 5% milk in standard tris base, saline tween (TBST) and then probed with a rabbit polyclonal antibody (Thermo Fisher Scientific, Waltham, MA) at 1:100 in blocking buffer overnight at 4C. For crygs (Thermo Fisher Scientific, Waltham, MA) detection, separate membranes were blocked in 5% BSA in TBST and then probed with a rabbit polyclonal antibody (Thermo Fisher Scientific, Waltham, MA) at 1:100 in blocking buffer overnight at 4C. The resulting bands were visualized with a secondary antibody Anti-rabbit IgG HRP-linked antibody (Thermo Fisher Scientific, Waltham, MA) at 1:1000 for 1 hour at room temperature. Membranes where then stripped and re-probed for the loading control actin or tubulin. An Anti-actin polyclonal antibody (Millipore Sigma, St. Louis, MO) made in mouse was diluted at 1:1000 in TBST and incubated with the membrane overnight at 4C. The resulting bands were visualized with a secondary antibody Anti-mouse IgG HRP-linked antibody (Cell Signaling Technology, Dancers, MA) at 1:1000 for 1 hour at room temperature. Protein densities for cryba and cryg2 bands were measured and normalized to the protein densities of the corresponding actin bands. Each sample was at least triplicated and averaged according to each gnotobiotic derivation.

### ATP Measurement and Gavage

For measuring total ATP concentration of whole larvae, 6/7dpf zebrafish were euthanized by tricane. Individual zebrafish were put into an Eppendorf tube, pipetting out as much excess liquid as possible and 100 µl of 95C distilled water was added to each fish. The tubes were immediately but on the 95C heat-block. All tubes of zebrafish were incubated at 95C for 20-30 minutes, mixing by pipette every 5 minutes. All sample carcass remnants were pellet by centrifugation at 14,000 rpm for 30 minutes at 4C. The ATP concentration of the supernatant was calculated by the ATP Determination Kit (Thermo Scientific, Rockford, IL) relative to a standard. The same protocol was used to measure the ATP concentration of dissected digestive systems of larvae but dissected tissue was boiled in 30 µl of 95C water and centrifugation was omitted. The ADP/ATP Ratio Assay Kit (Bioluminescent) (Abcam, Cambridge, MA) was used to measure the average ratio of ADP to ATP of larvae within dissected digestive systems from 2 zebrafish per sample using the same tissue preparation with 95C water as described above.

To measure ATP turnover in the larval intestine, 6dpf zebrafish were gavaged with 1 or 10mM of ATP (Millipore Sigma, St. Louis, MO), dissolved in embryo media, directly into the proximal intestine as previously described (Cocchiaro and Rawls, 2013). Approximately 20-40 minutes post gavage, the digestive systems of zebrafish were dissected and processed for measuring ATP concentration as described above.

### Statistical Analysis

Data from *in vivo* experiments are displayed as boxplots with the data median as the line within the box, the top and bottom of the box representing the upper and lower quartile, the whiskers representing the min and max values, and the ‘+’ symbol representing the mean. A Students’ Ttest was used to gauge statistical differences of means between CVZ and GF groups in non-sequencing experiments. For all statistical tests, p<0.05 is considered significant.

### Accession of Data

All data will be readily available at the time of publication

## References

Adolph, T. E., Mayr, L., Grabherr, F., Schwärzler, J., and Tilg, H. (2019). Pancreas-Microbiota Cross Talk in Health and Disease. Annual Review of Nutrition, 39, 249–266. https://doi.org/10.1146/annurev-nutr-082018-124306

Arentsen, T., Qian, Y., Gkotzis, S., Femenia, T., Wang, T., Udekwu, K., and Diaz Heijtz, R. (2016). The bacterial peptidoglycan-sensing molecule Pglyrp2 modulates brain development and behavior. https://doi.org/10.1038/mp.2016.182

Atarashi, K., Nishimura, J., Shima, T., Umesaki, Y., Yamamoto, M., Onoue, M., and Takeda, K. (2008). ATP drives lamina propria TH17 cell differentiation. Nature 2008 455:7214, 455(7214), 808–812. https://doi.org/10.1038/nature07240

Bao, Y., Ledderose, C., Seier, T., Graf, A. F., Brix, B., Chong, E., and Junger, W. G. (2014). Mitochondria regulate Neutrophil activation by generating ATP for Autocrine Purinergic signaling. Journal of Biological Chemistry, 289(39), 26794–26803. https://doi.org/10.1074/jbc.M114.572495

Bates, J. M., Mittge, E., Kuhlman, J., Baden, K. N., Cheesman, S. E., and Guillemin, K. (2006). Distinct signals from the microbiota promote different aspects of zebrafish gut differentiation. Developmental Biology. https://doi.org/10.1016/j.ydbio.2006.05.006

Biemar, F., Argenton, F., Schmidtke, R., Epperlein, S., Peers, B., and Driever, W. (2001). Pancreas development in zebrafish: early dispersed appearance of endocrine hormone expressing cells and their convergence to form the definitive islet. Developmental Biology, 230(2), 189–203. https://doi.org/10.1006/dbio.2000.0103

Bott, C. J., and Winckler, B. (2020). Intermediate filaments in developing neurons: Beyond structure. https://doi.org/10.1002/cm.21597

Bruckner, J. J., Stednitz, S. J., Grice, M. Z., Larsch, J., Tallafuss, A., Washbourne, P., and Eisen, J. S. (2020). The microbiota promotes social behavior by neuro-immune modulation of neurite complexity. BioRxiv, 2020.05.01.071373. https://doi.org/10.1101/2020.05.01.071373

Cheesman, S. E., Neal, J. T., Mittge, E., Seredick, B. M., and Guillemin, K. (2011). Epithelial cell proliferation in the developing zebrafish intestine is regulated by the Wnt pathway and microbial signaling via Myd88.https://doi.org/10.1073/pnas.1000072107

Cocchiaro, J. L., and Rawls, J. F. (2013). Microgavage of zebrafish larvae. Journal of Visualized Experiments, (72). https://doi.org/10.3791/4434

Cui, X et al. (2020). Heat shock factor 4 regulates lysosome activity by modulating the αB-crystallin-ATP6V1A-mTOR complex in ocular lens. Biochimica et Biophysica Acta (BBA) - General Subjects, 1864(3), 129496. https://doi.org/10.1016/J.BBAGEN.2019.129496

DiIorio, P. J., Moss, J. B., Sbrogna, J. L., Karlstrom, R. O., and Moss, L. G. (2002). Sonic hedgehog Is Required Early in Pancreatic Islet Development. Developmental Biology, 244(1), 75–84. https://doi.org/10.1006/DBIO.2002.0573

Dong, P. D. S., Provost, E., Leach, S. D., and Stainier, D. Y. R. (2008). Graded levels of Ptf1a differentially regulate endocrine and exocrine fates in the developing pancreas. Genes and Development, 22(11), 1445–1450. https://doi.org/10.1101/gad.1663208

Donohoe, D. R., Garge, N., Zhang, X., Sun, W., O’Connell, T. M., Bunger, M. K., and Bultman, S. J. (2011). The microbiome and butyrate regulate energy metabolism and autophagy in the mammalian colon. Cell Metabolism, 13(5), 517–526. https://doi.org/10.1016/J.CMET.2011.02.018/ATTACHMENT/47419042-0759-4B45-AADD-3942F67E2AE4/MMC3.XLS

Farnsworth, D. R., Posner, M., and Miller, A. C. (2021). Single cell transcriptomics of the developing zebrafish lens and identification of putative controllers of lens development. Experimental Eye Research, v206. https://doi.org/10.1016/J.EXER.2021.108535

Farnsworth, D. R., Saunders, L. M., and Miller, A. C. (2019). A single-cell transcriptome atlas for zebrafish development. Developmental Biology. https://doi.org/10.1016/j.ydbio.2019.11.008

Field, H. A., Si Dong, P. D., Beis, D., and Stainier, D. Y. R. (2003). Formation of the digestive system in zebrafish. ii. pancreas morphogenesis☆. Developmental Biology, 261(1), 197– 208. https://doi.org/10.1016/S0012-1606(03)00308-7

Frost, F et al. (2019). Impaired Exocrine Pancreatic Function Associates With Changes in Intestinal Microbiota Composition and Diversity. Gastroenterology, 156(4), 1010–1015. https://doi.org/10.1053/j.gastro.2018.10.047

Fu, X., Ezemaduka, A. N., Lu, X., and Chang, Z. (2021). The Caenorhabditis elegans 12-kDa small heat shock proteins with little in vitro chaperone activity play crucial roles for its dauer formation, longevity, and reproduction. Protein Science, 30(10), 2170–2182. https://doi.org/10.1002/PRO.4160

Garbett, K. A., Hsiao, E. Y., Kálmán, S., Patterson, P. H., and Mirnics, K. (2012). Effects of maternal immune activation on gene expression patterns in the fetal brain. Translational Psychiatry, 2. https://doi.org/10.1038/tp.2012.24

Gnainsky, Y et al. (2021). Systemic Regulation of Host Energy and Oogenesis by Microbiome-Derived Mitochondrial Coenzymes. Cell Reports, 34(1). https://doi.org/10.1016/j.celrep.2020.108583

Godinho, L et al. (2005). Targeting of amacrine cell neurites to appropriate synaptic laminae in the developing zebrafish retina. Development, 132(22), 5069–5079. https://doi.org/10.1242/dev.02075

Groh, K. J., and J-F Suter M., (2015). Stressor-induced proteome alterations in zebrafish: A meta-analysis of response patterns. Aquatic Toxicology, 159, 1–12. https://doi.org/10.1016/j.aquatox.2014.11.013

Heppert, J. K., Davison, J. M., Kelly, C., Mercado, G. P., Lickwar, C. R., and Rawls, J. F. (2021, January 1). Transcriptional programmes underlying cellular identity and microbial responsiveness in the intestinal epithelium. Nature Reviews Gastroenterology and Hepatology, Vol. 18, pp. 7–23. https://doi.org/10.1038/s41575-020-00357-6

Holm En Larsson, J.M., Karlsson, H., Crespo, J. G., Johansson, M. E. V, Eklund, L., Sjövall, H., and Hansson, G. C. (2011). Altered O-glycosylation Profile of MUC2 Mucin Occurs in Active Ulcerative Colitis and Is Associated with Increased Inflammation. https://doi.org/10.1002/ibd.21625

Hotamisligil, G. S. (2017). Inflammation, metaflammation and immunometabolic disorders. Nature 2017 542:7640, 542(7640), 177–185. https://doi.org/10.1038/nature21363

Howe, D. G. et al. (2013). ZFIN, the Zebrafish Model Organism Database: Increased support for mutants and transgenics. Nucleic Acids Research, 41(D1), 854–860. https://doi.org/10.1093/nar/gks938

Hsiao, E. Y. et al. (2013). Microbiota modulate behavioral and physiological abnormalities associated with neurodevelopmental disorders. Cell, 155(7), 1451–1463. https://doi.org/10.1016/j.cell.2013.11.024

Huang, L., Otrokocsi, L., and Sperlágh, B. (2019, September 1). Role of P2 receptors in normal brain development and in neurodevelopmental psychiatric disorders. Brain Research Bulletin, Vol. 151, pp. 55–64. https://doi.org/10.1016/j.brainresbull.2019.01.030

Iwanami, N., Lawir, D. F., Sikora, K., O’Meara, C., Takeshita, K., Schorpp, M., and Boehm, T. (2020). Transgenerational inheritance of impaired larval T cell development in zebrafish. Nature Communications, 11(1). https://doi.org/10.1038/S41467-020-18289-9

Jameson, K. G., Olson, C. A., Kazmi, S. A., and Hsiao, E. Y. (2020). Toward Understanding Microbiome-Neuronal Signaling. Molecular Cell, 78(4), 577–583. https://doi.org/10.1016/J.MOLCEL.2020.03.006

Jevtov, I., Samuelsson, T., Yao, G., Amsterdam, A., and Ribbeck, K. (2014). Zebrafish as a model to study live mucus physiology OPEN. https://doi.org/10.1038/srep06653

Kang, B. et al. (2019). Commensal microbiota drive the functional diversification of colon macrophages. Mucosal Immunology. https://doi.org/10.1038/s41385-019-0228-3

Kanther, M. et al. (2014). Commensal microbiota stimulate systemic neutrophil migration through induction of Serum amyloid A. Cellular Microbiology, 16(7), 1053–1067. https://doi.org/10.1111/CMI.12257/SUPPINFO

Khodajou-Masouleh, H., Shahangian, S. S., Attar, F., H. Sajedi R.,, and Rasti, B. (2021). Characteristics, dynamics and mechanisms of actions of some major stress-induced biomacromolecules; addressing Artemia as an excellent biological model. Journal of Biomolecular Structure & Dynamics, 39(15), 5619–5637. https://doi.org/10.1080/07391102.2020.1796793

King, A. M., and Macrae, T. H. (2015). Insect heat shock proteins during stress and diapause. Annual Review of Entomology, 60, 59–75. https://doi.org/10.1146/ANNUREV-ENTO-011613-162107

Kurup, N., Li, Y., Goncharov, A., and Jin, Y. (2018). Intermediate filament accumulation can stabilize microtubules in Caenorhabditis elegans motor neurons. Proceedings of the National Academy of Sciences of the United States of America, 115(12), 3114–3119. https://doi.org/10.1073/PNAS.1721930115/VIDEO-4

Lickwar, C. R. et al. (2017). Genomic dissection of conserved transcriptional regulation in intestinal epithelial cells. https://doi.org/https://doi.org/10.1371/journal.pbio.2002054

MacVicar, B. A., Wicki-Stordeur, L., and Bernier, L. P. (2017, May 22). The cost of communication in the brain. ELife, Vol. 6. https://doi.org/10.7554/eLife.27894

Margulis, L. (1976). Genetic and evolutionary consequences of symbiosis. Experimental Parasitology, 39(2), 277–349. https://doi.org/10.1016/0014-4894(76)90127-2

Mcfall-Ngai, M. J. (2014). MI68CH10-McFall-Ngai The Importance of Microbes in Animal Development: Lessons from the Squid-Vibrio Symbiosis. https://doi.org/10.1146/annurev-micro-091313-103654

Meisel, J. S. et al. (2018). Commensal microbiota modulate gene expression in the skin. https://doi.org/10.1186/s40168-018-0404-9

Melancon, E. et al. (2017). Best practices for germ-free derivation and gnotobiotic zebrafish husbandry. https://doi.org/10.1016/bs.mcb.2016.11.005

Mishra, S., Wu, S. Y., Fuller, A. W., Wang, Z., Rose, K. L., Schey, K. L., and Mchaourab, H. S. (2018). Loss of αb-crystallin function in zebrafish reveals critical roles in the development of the lens and stress resistance of the heart. Journal of Biological Chemistry, 293(2), 740– 753. https://doi.org/10.1074/JBC.M117.808634/ATTACHMENT/A574346B-3530-4F3F-8470-27770850FABD/MMC2.PDF

Murdoch, C. C., and Rawls, J. F. (2019, September 6). Commensal Microbiota Regulate Vertebrate Innate Immunity-Insights From the Zebrafish. Frontiers in Immunology, Vol. 10, p. 2100. https://doi.org/10.3389/fimmu.2019.02100

Nishiyama, H. et al. (2018). Supplementation of pancreatic digestive enzymes alters the composition of intestinal microbiota in mice. Biochemical and Biophysical Research Communications, 495(1), 273–279. https://doi.org/10.1016/j.bbrc.2017.10.130

Park, J. et al. (2019). Lysosome-Rich Enterocytes Mediate Protein Absorption in the Vertebrate Gut Article Lysosome-Rich Enterocytes Mediate Protein Absorption in the Vertebrate Gut. Developmental Cell, 1–14. https://doi.org/10.1016/j.devcel.2019.08.001

Pauls, S., Zecchin, E., Tiso, N., Bortolussi, M., and Argenton, F. (2007). Function and regulation of zebrafish nkx2.2a during development of pancreatic islet and ducts. Developmental Biology, 304(2), 875–890. https://doi.org/10.1016/j.ydbio.2007.01.024

Phadte, A. S., Sluzala, Z. B., and Fort, P. E. (2021). Therapeutic Potential of α-Crystallins in Retinal Neurodegenerative Diseases. Antioxidants 2021, Vol. 10, Page 1001, 10(7), 1001. https://doi.org/10.3390/ANTIOX10071001

Pietzner, M. et al. (2021). Exocrine Pancreatic Function Modulates Plasma Metabolites Through Changes in Gut Microbiota Composition. The Journal of Clinical Endocrinology & Metabolism, XX, 1–9. https://doi.org/10.1210/clinem/dgaa961

Piri, N., Kwong, J. M. K., and Caprioli, J. (2013). Crystallins in Retinal Ganglion Cell Survival and Regeneration. Molecular Neurobiology, 48(3), 819–828. https://doi.org/10.1007/s12035-013-8470-2

Pronovost, G. N., and Hsiao, E. Y. (2019). Perinatal Interactions between the Microbiome, Immunity, and Neurodevelopment. Immunity, 50(1), 18–36. https://doi.org/10.1016/j.immuni.2018.11.016

Rawls, J. F., Samuel, B. S., and Gordon, J. I. (2004). Gnotobiotic zebrafish reveal evolutionarily conserved responses to the gut microbiota. Retrieved from www.pnas.orgcgidoi10.1073pnas.0400706101

Rolig, A. S., Parthasarathy, R., Burns, A. R., Bohannan, B. J. M., and Guillemin, K. (2015). Individual members of the microbiota disproportionately modulate host innate immune responses. Cell Host and Microbe, 18(5), 613–620. https://doi.org/10.1016/j.chom.2015.10.009

Sampson, T. R., and Mazmanian, S. K. (2015). Control of brain development, function, and behavior by the microbiome. Cell Host and Microbe. https://doi.org/10.1016/j.chom.2015.04.011

Semova, I., Carten, J. D., Stombaugh, J., MacKey, L. C., Knight, R., Farber, S. A., and Rawls, J. F. (2012). Microbiota regulate intestinal absorption and metabolism of fatty acids in the zebrafish. Cell Host and Microbe, 12(3), 277–288. https://doi.org/10.1016/j.chom.2012.08.003

Sharma, P. V., and Thaiss, C. A. (2020). Host-Microbiome Interactions in the Era of Single-Cell Biology. Frontiers in Cellular and Infection Microbiology, 10, 536. https://doi.org/10.3389/FCIMB.2020.569070/BIBTEX

Shepherd, I., and Eisen, J. (n.d.). Development of the Zebrafish Enteric Nervous System. https://doi.org/10.1016/B978-0-12-387036-0.00006-2

Shih, L. J., Lu, Y. F., Chen, Y. H., Lin, C. C., Chen, J. A., and Hwang, S. P. L. (2007). Characterization of the agr2 gene, a homologue of X. laevis anterior gradient 2, from the zebrafish, Danio rerio. Gene Expression Patterns, 7(4), 452–460. https://doi.org/10.1016/j.modgep.2006.11.003

Singh, S. P. et al. (2022). STEM CELLS AND REGENERATION A single-cell atlas of de novo β-cell regeneration reveals the contribution of hybrid β/δ-cells to diabetes recovery in zebrafish. https://doi.org/10.1242/dev.199853

Slingsby, C., and Wistow, G. J. (2014). Functions of crystallins in and out of lens: Roles in elongated and post-mitotic cells. Progress in Biophysics and Molecular Biology, 115(1), 52–67. https://doi.org/10.1016/J.PBIOMOLBIO.2014.02.006

Spencer, N. J., and Hu, H. (2020). Enteric nervous system: sensory transduction, neural circuits and gastrointestinal motility. Nature Reviews. Gastroenterology & Hepatology, 17(6), 338– 351. https://doi.org/10.1038/S41575-020-0271-2

Sprague-Piercy, M. A., Rocha, M. A., Kwok, A. O., and Martin, R. W. (2021). α-Crystallins in the Vertebrate Eye Lens: Complex Oligomers and Molecular Chaperones. Annual Review of Physical Chemistry, 72, 143–163. https://doi.org/10.1146/ANNUREV-PHYSCHEM-090419-121428

Stuart, T. et al. (2018). Comprehensive integration of single cell data. BioRxiv, 460147. https://doi.org/10.1101/460147

Taylor, C. R., Montagne, W. A., Eisen, J. S., and Ganz, J. (2016). Molecular fingerprinting delineates progenitor populations in the developing zebrafish enteric nervous system. Developmental Dynamics, 245(11), 1081–1096. https://doi.org/10.1002/dvdy.24438

Troll, J. V. et al. (2018). Microbiota promote secretory cell determination in the intestinal epithelium by modulating host Notch signaling. https://doi.org/10.1242/dev.155317

Udayangani, R. M. C., Dananjaya, S. H. S., Nikapitiya, C., Heo, G. J., Lee, J., and De Zoysa, M. (2017). Metagenomics analysis of gut microbiota and immune modulation in zebrafish (Danio rerio) fed chitosan silver nanocomposites. Fish and Shellfish Immunology, 66, 173– 184. https://doi.org/10.1016/j.fsi.2017.05.018

Valapala, M. et al. (2016). Modulation of V-ATPase by βA3/A1-Crystallin in Retinal Pigment Epithelial Cells. Advances in Experimental Medicine and Biology, 854, 779–784. https://doi.org/10.1007/978-3-319-17121-0_104

Verderio, C., and Matteoli, M. (2011, January 7). ATP in neuron-glia bidirectional signalling. Brain Research Reviews, Vol. 66, pp. 106–114. https://doi.org/10.1016/j.brainresrev.2010.04.007

Wan, H. et al. (2006). Analyses of pancreas development by generation of gfp transgenic zebrafish using an exocrine pancreas-specific elastaseA gene promoter. Experimental Cell Research, 312(9), 1526–1539. https://doi.org/10.1016/J.YEXCR.2006.01.016

Wen, J. et al.(2021). Fxr signaling and microbial metabolism of bile salts in the zebrafish intestine. Science Advances, 7(30). https://doi.org/10.1126/SCIADV.ABG1371

Westerfield, M. (2007) The Zebrafish Book. A Guide for the Laboratory Use of Zebrafish (Danio rerio), 5th Edition. University of Oregon Press, Eugene. https://zfin.org/ZDB-PUB-101222-53

Wiles, T. J., and Guillemin, K. (2020, April 1). Patterns of partnership: surveillance and mimicry in host-microbiota mutualisms. Current Opinion in Microbiology, Vol. 54, pp. 87–94. https://doi.org/10.1016/j.mib.2020.01.012

Willms, R. J., Jones, L. O., Hocking, J. C., and Foley, E. (2022). A cell atlas of microbe-responsive processes in the zebrafish intestine. Cell Reports, 38(5), 110311. https://doi.org/10.1016/J.CELREP.2022.110311

Zhu, X. H. et al. (2012). Quantitative imaging of energy expenditure in human brain. NeuroImage, 60(4), 2107–2117. https://doi.org/10.1016/j.neuroimage.2012.02.013

Zhu, Z., and Reiser, G. (2018). The small heat shock proteins, especially HspB4 and HspB5 are promising protectants in neurodegenerative diseases. Neurochemistry International, 115, 69–79. https://doi.org/10.1016/J.NEUINT.2018.02.006

Zigler, J. S., and Sinha, D. (2015). βA3/A1-crystallin: more than a lens protein. Progress in Retinal and Eye Research, 44, 62–85. https://doi.org/10.1016/J.PRETEYERES.2014.11.002

