## Supplementary figures and images for "GLOBAL HOST RESPONSES TO THE MICROBIOTA AT SINGLE CELL RESOLUTION IN GNOTOBIOTIC ZEBRAFISH"

### Supplemental Figure 1

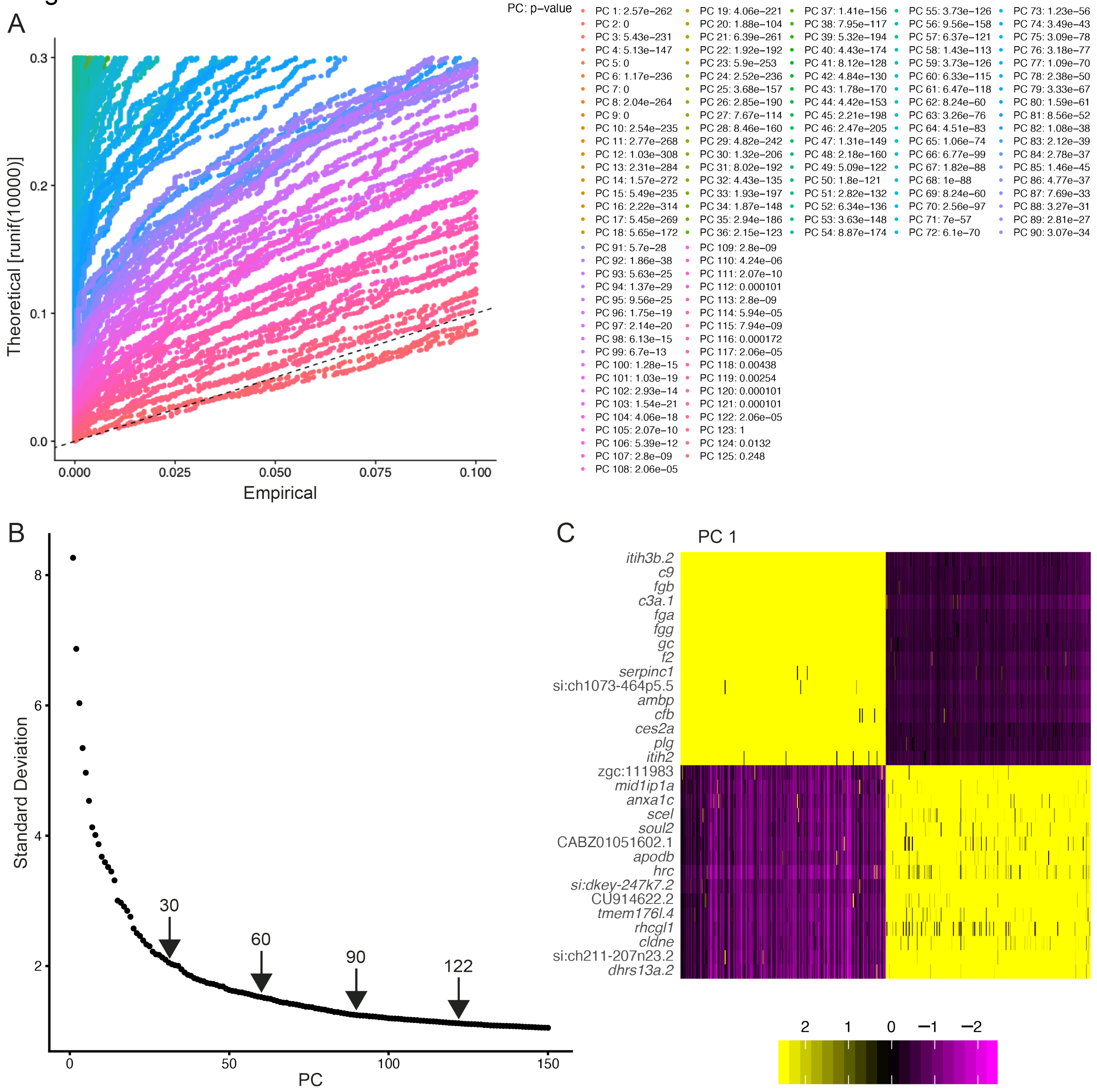

### Supplemental Figure 2

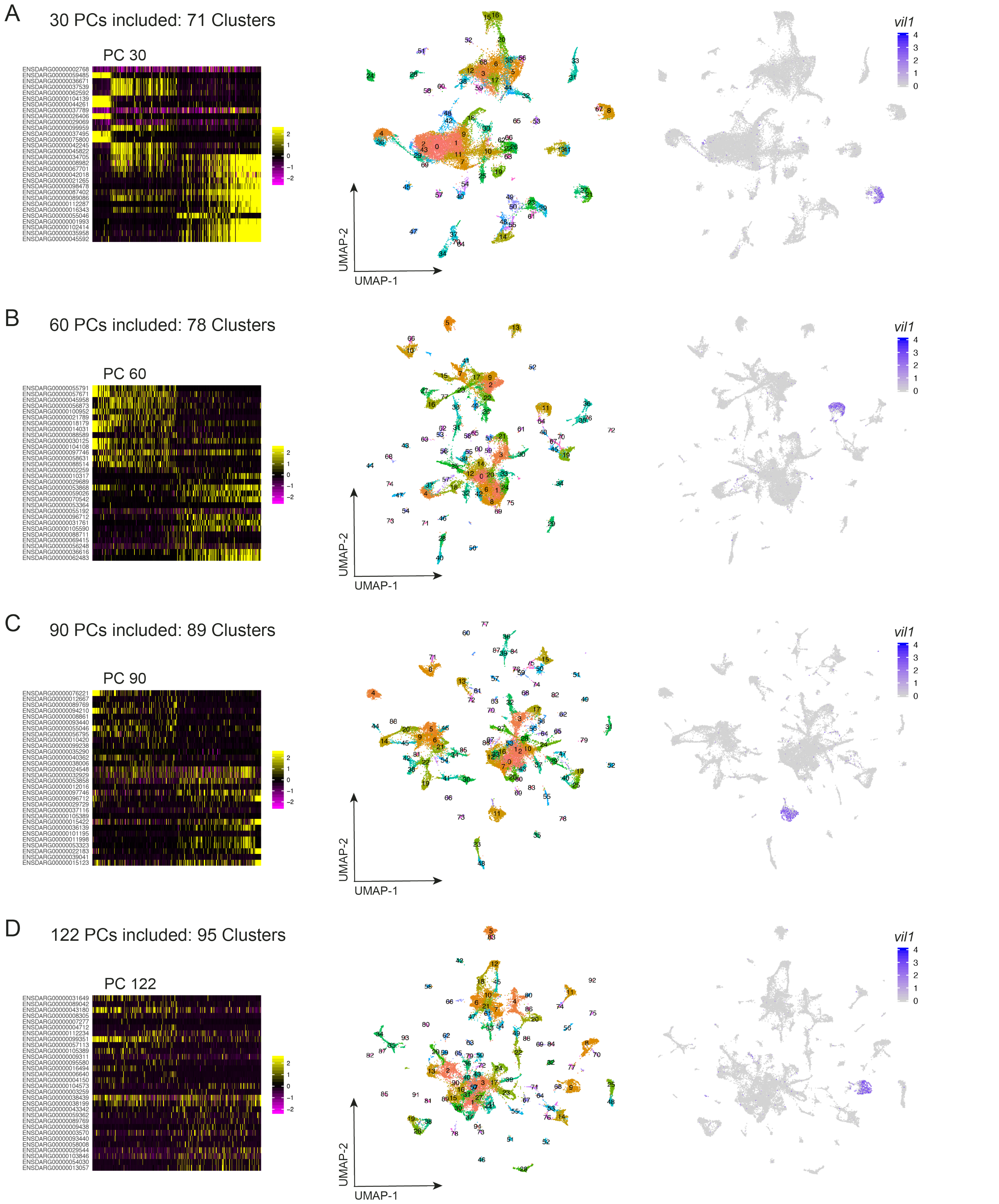

### Supplemental Figure 3

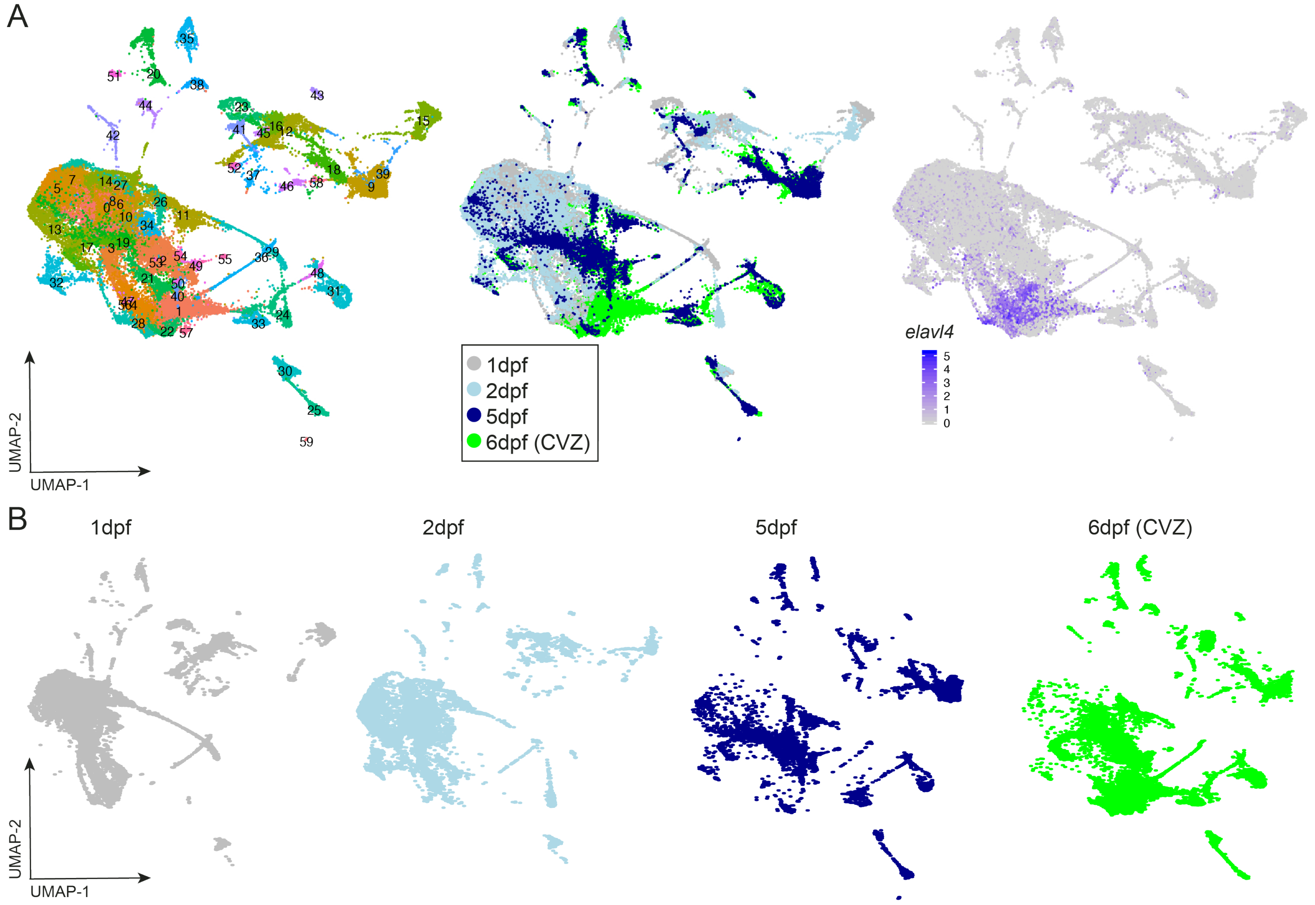

### Supplemental Figure 4

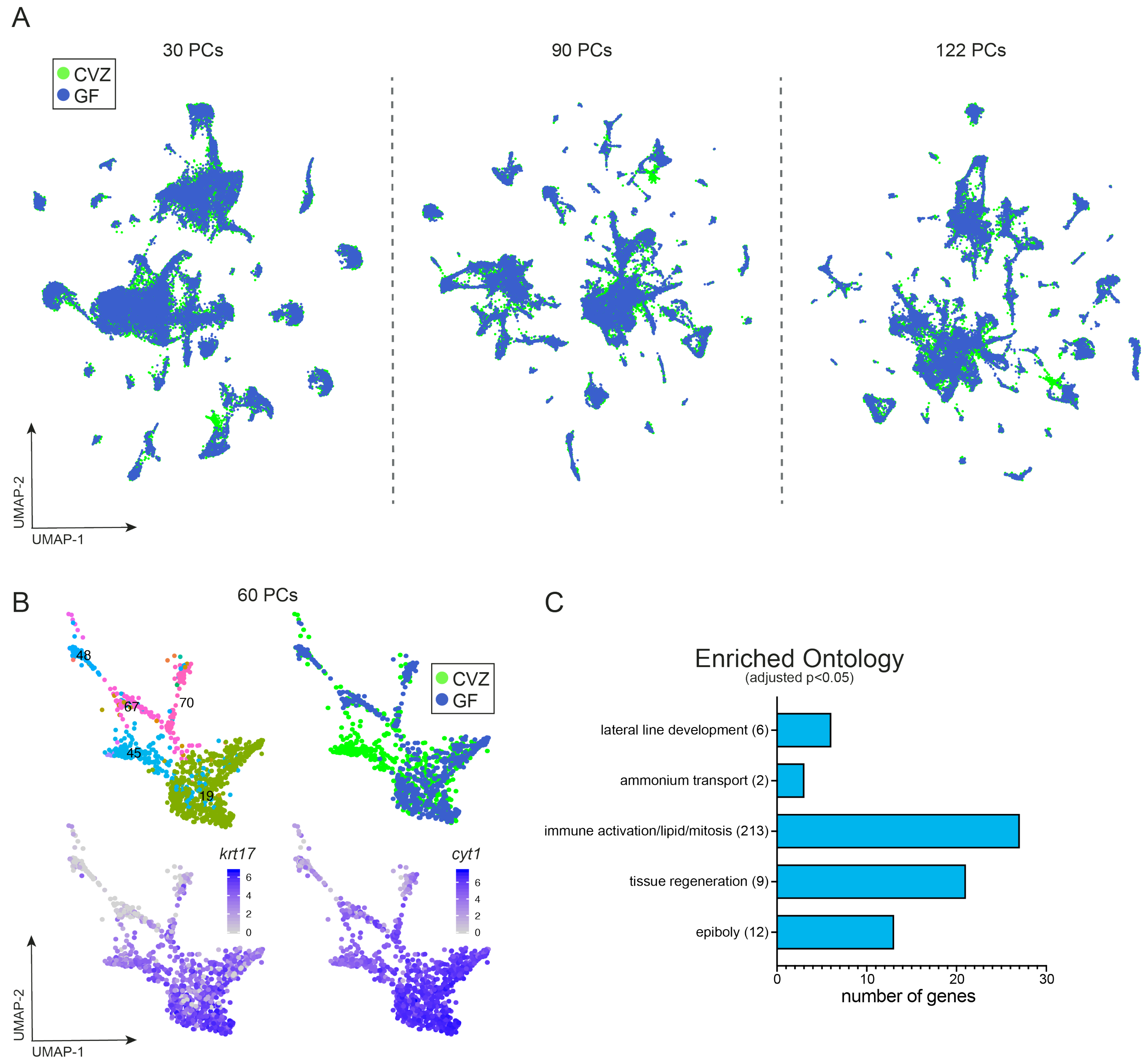

### Supplemental Figure 5

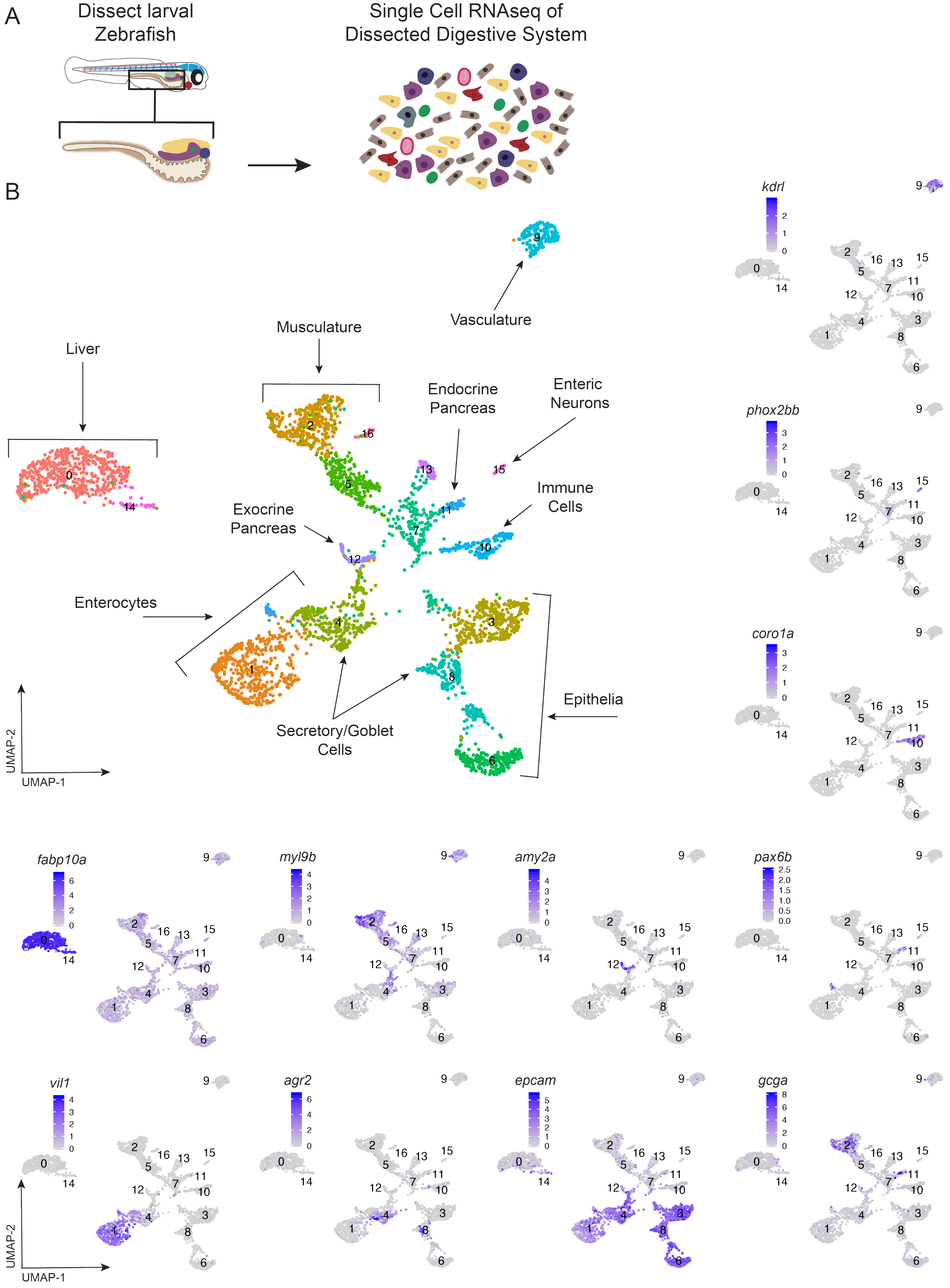

### Supplemental Figure 7

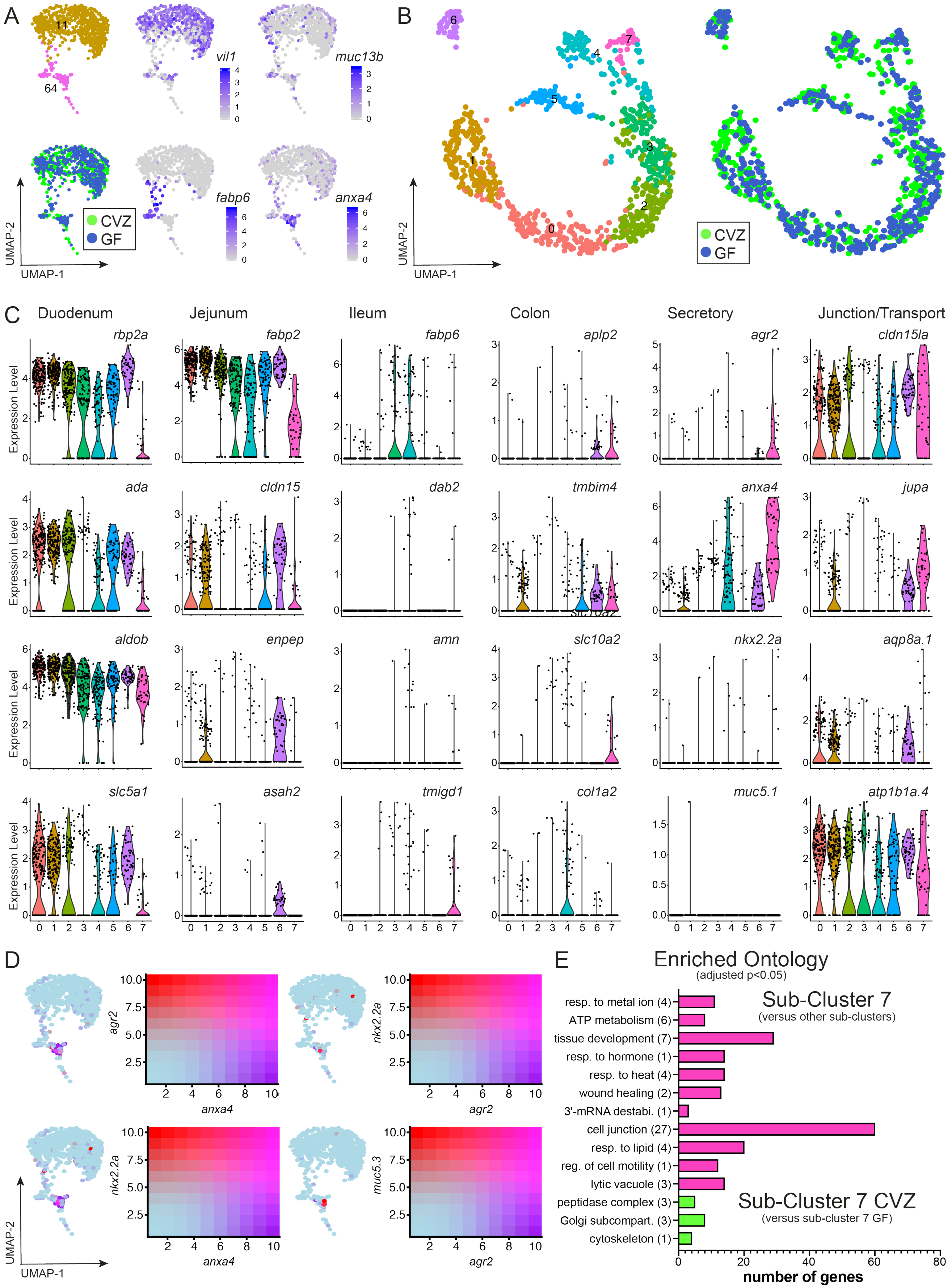

### Supplemental Figure 8

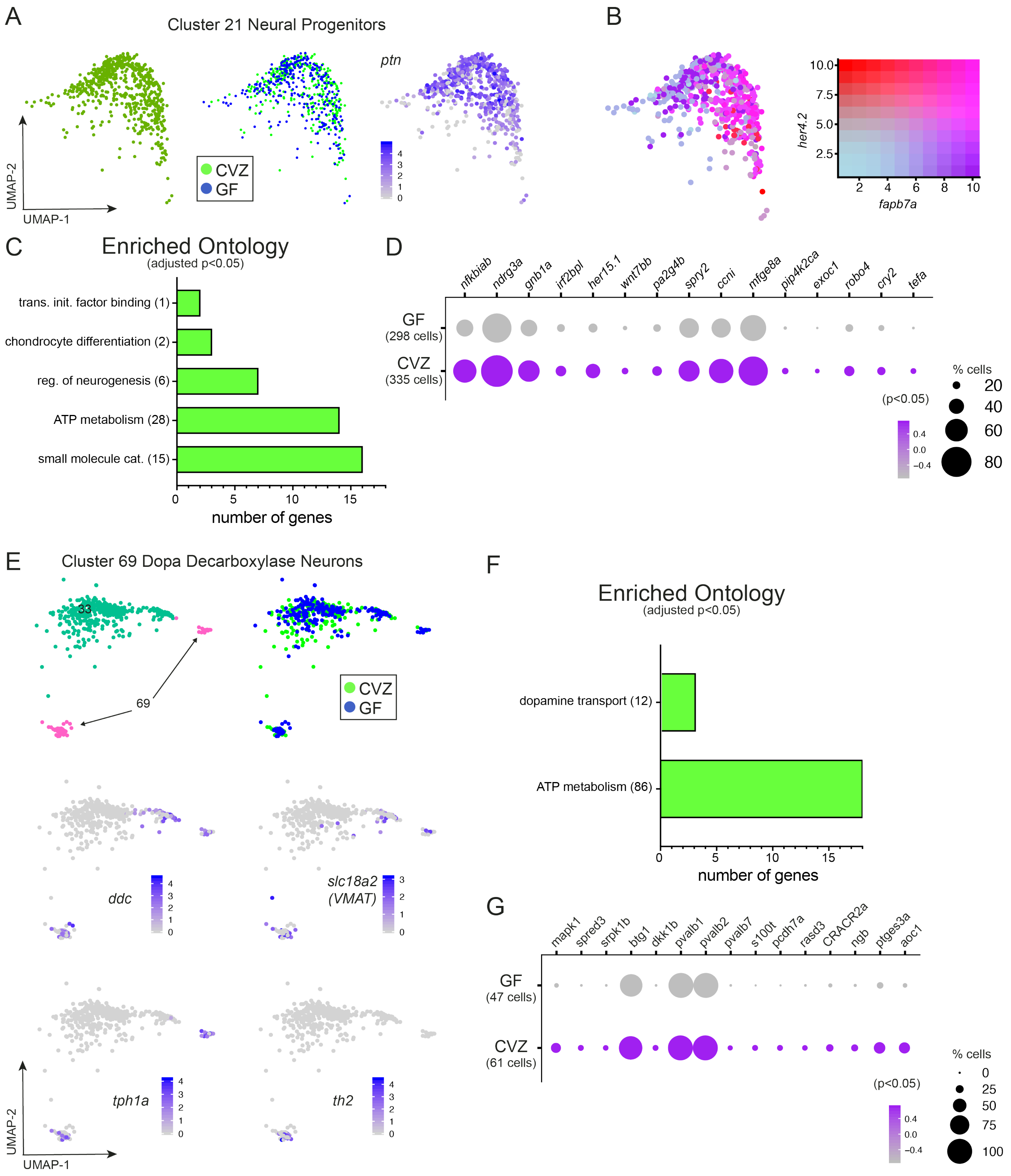

### Supplemental Figure 9

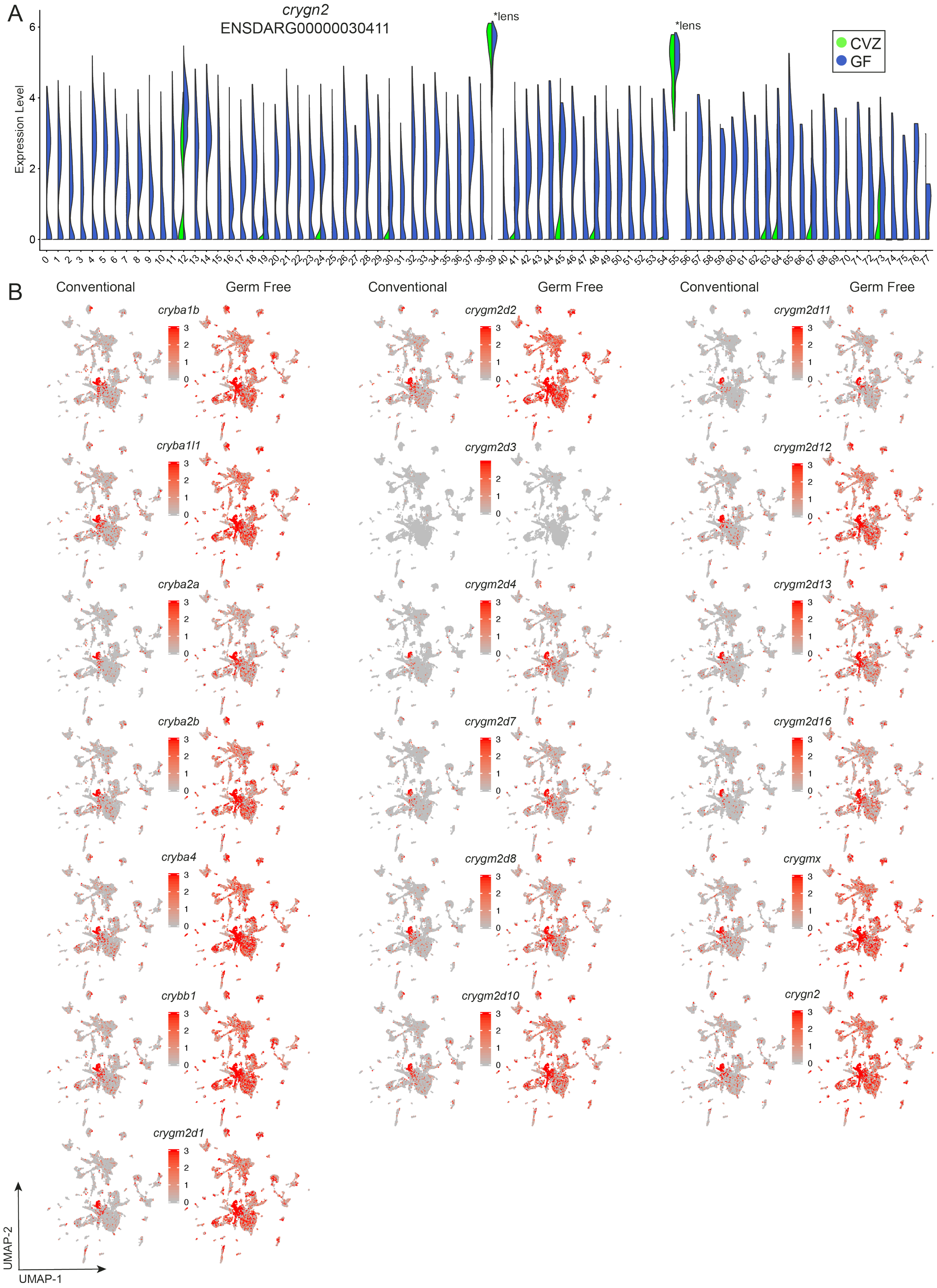

### Supplemental Figure 10

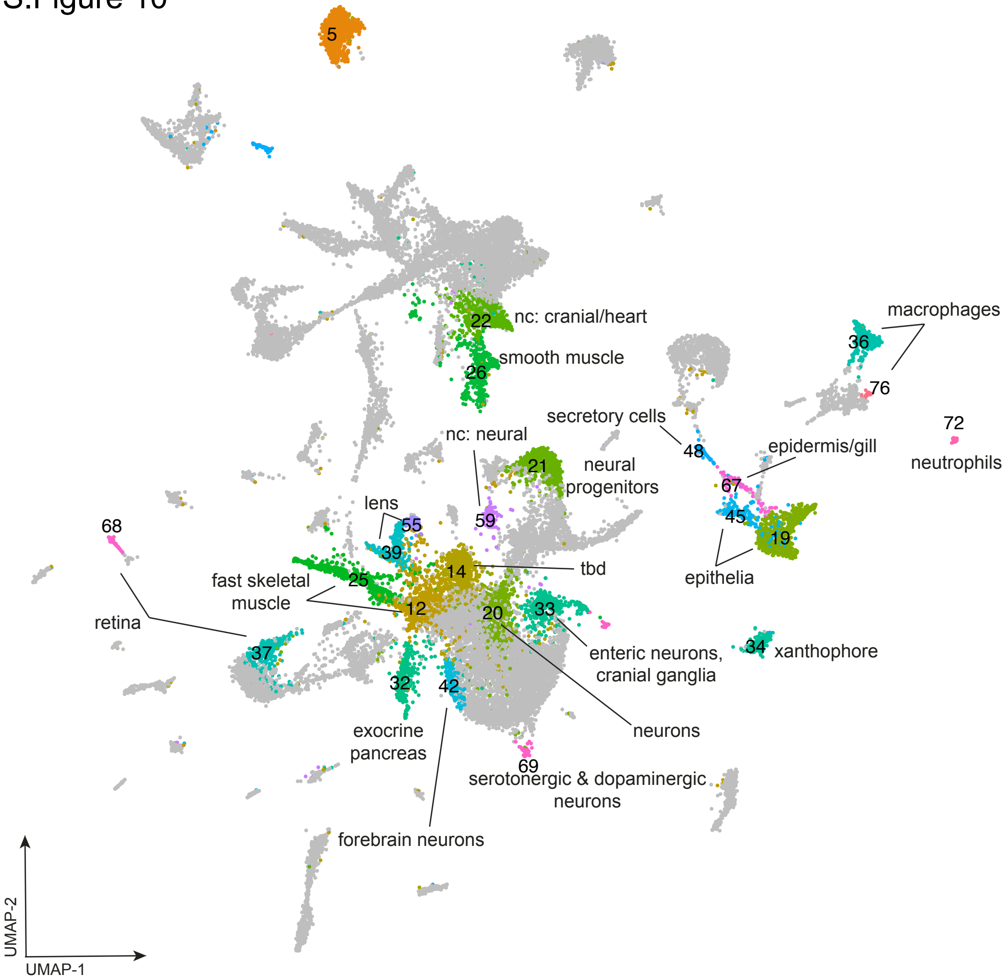

### Supplemental Figure 11

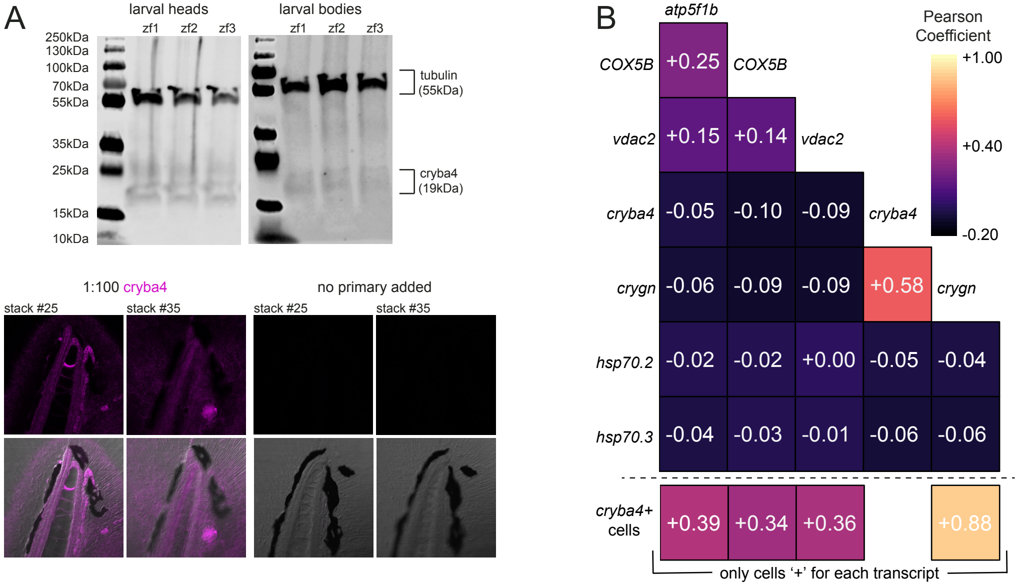

### Supplemental Figure 12

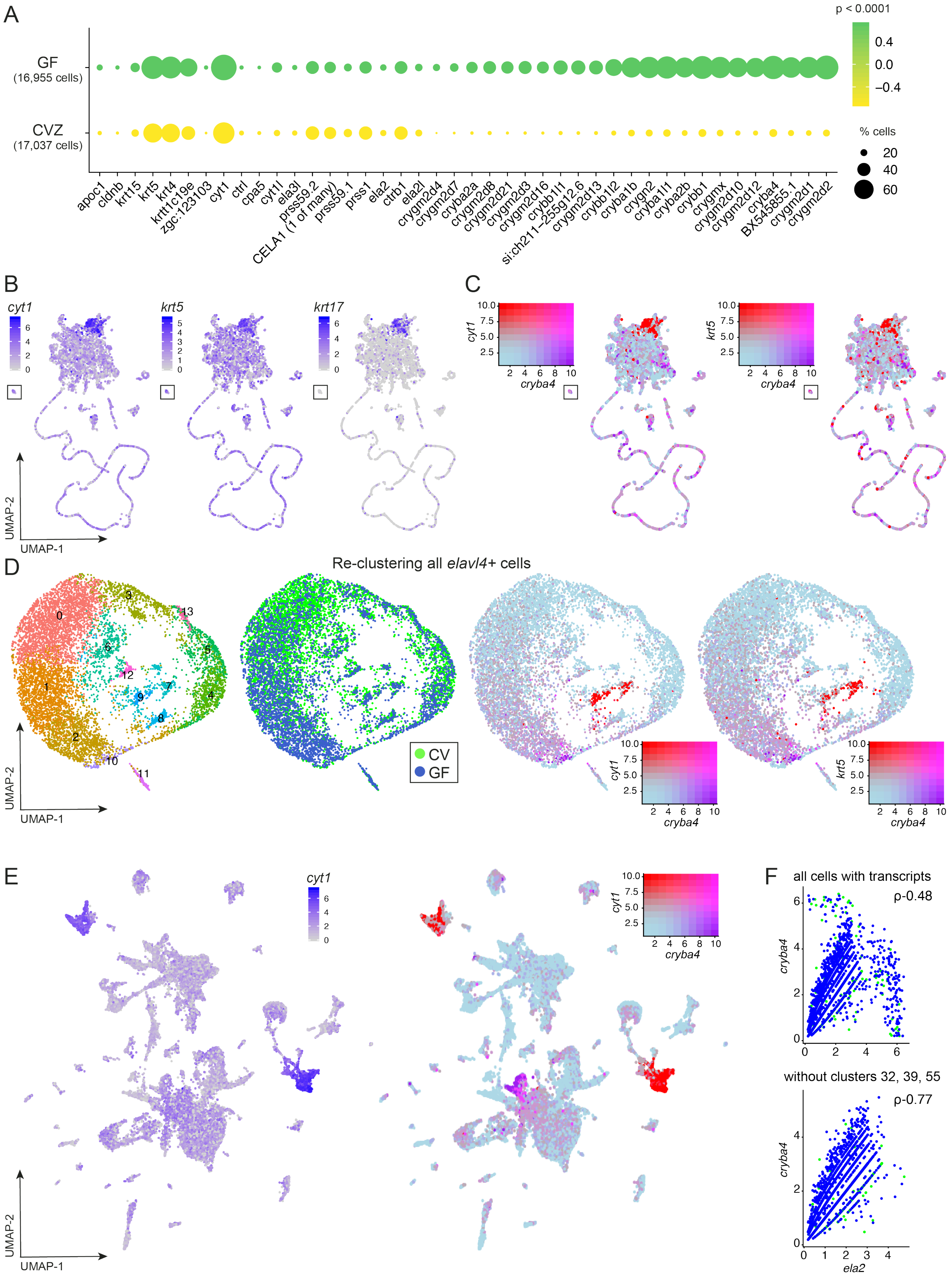
