## Supplemental Figure 6 for "GLOBAL HOST RESPONSES TO THE MICROBIOTA AT SINGLE CELL RESOLUTION IN GNOTOBIOTIC ZEBRAFISH"

S.Figure 6

### Comparing Enriched Gene Expression in CV Cells

(Rawls et al., 2004 Microarray on Dissected Guts versus Single Cell Clusters)

Similar Trend

Opposite Trend

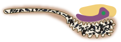

| Rawls et al 2004 DEGs | Cluster 11 & 64 **Enterocytes** | Cluster 28 & 40 **Liver** | Cluster 48 & 61 **Secretory Cells** | Cluster 33 **Enteric Neurons** | Cluster 32 **Exocrine Pancreas** | Cluster 72 **Neutrophils** | Cluster 36 & 76 **Macrophages** |
|---|---|---|---|---|---|---|---|
| **ANXA2** | anxa4 | anxa4 | anxa1a | anxa1a | anxa1a | anxa4 | anxa1a, anxa11b |
| **AP1S1** | | | | | ap2m1a | | ap1s2, ap2s1, ap3s2 |
| **Apob** | apoa4b.1, apoa4b.2 (2x), apoc2, | apoc1, apom | apoa1b, apoa2 | apoa1b, apoa2, apoe | apoc1, apoda.2 | apoa1b, apoa2, apoc1, apom | |
| | apoa1b, apoa2 | | | | | | |
| **ARP2** | arpc1a | | | | arpc2 | | |
| **C3** | | c3a.3 | | | | | |
| **C4** | | c4b | | | | | |
| **Calr$^g$** | calr | | calr3a | calr | calr | | |
| **CBX1** | | cbx7a | | | cbx1a, cbx1b, cbx7a | | |
| **CORO1C** | | | | | | coro1a | |
| **Dlc2** | | dynll2b | | | | | |
| **DNAJB11** | | dnajc5aa | dnajc5aa | dnaja2a, dnajc7, dnajc8 | dnaja2b, dnajc5aa | | |
| | | | dnajb12a | | | | |
| **Gpx2** | gpx1b | gpx1a | gpx1a, gpx1b | | | | gpx1a |
| | | | gpx4a | | | | |
| **HMGA1** | hmgb1b | hmgb1a, hmgb1b | hmga1a, hmgb1b | | hmgb1a, hmgb1b | | |
| **HMGN2** | | hmgn7 | | | hmgn2, hmgn6, hmgn7 | | |
| **HSPD1$^h$** | hsp10.1, hsp70.2, hsp90b1, hspa5 | hsp70.2, hsp90aa1.2 | hsp90b1 | hsp70l | hsp70l, hsp90ab1 | hsp90ab1 | hsp70.2, hsp70.3 |
| **HSPD1$^h$** | | hspd1 | hspd1 | | | | |
| **IF2** | eif2s1b, eif3ha | eif1b, eif4a1b, eif4bb | | | | | |
| | eif3c, eif3d, eif3ja, eif3m, eif4abp2 | | | | | | |
| **Ifit1** | ifi45, ifi46 | | ifi46 | | | | |
| **KPNA2** | | | kpna3 | | | | |
| **LSM6** | | | lsm5, lsm6 | | | | |
| **MAPRE1$^g$** | map1ab | | | | map1aa, map1lc3b, map4l | mapre1a | |
| **Mcm5** | | | mcm5 | | | | |
| **MFAP4** | | | | | | | MFAP4 (1 of many) |
| **MSN** | msna | | | | | | |
| **NUCKS** | | nucks1a | | | nucks1a | | |
| **PABPC1** | | | pabpc1b | | pabpc1a | | |
| **Pcna** | | | pcna | | | | |
| **PFDN2** | | pfdn1 | | | pfdn1 | pfdn6 | |
| **PHB** | phb | | | | | | |
| **PPP1R3B** | | | ppp1r7 | | ppp2cb, ppp2r1ba | | ppp2cb |
| **PPP4R2** | ppp3r1b | | | | | | |
| **Psma5** | psma1, psma3, psma6a | | psma4, psma8 | | psma3, psma4, psma5 | psma3 | psma4, psma5, psma6a, psma6l |
| **Psmb3** | | | psmb4 | psmb1, psmb6 | | psmb1 | psmb6 |
| **Psmd12** | psmd2, psmd14 | psmd6 | psmc6 | | psmd1, psmd3 | psmd7, psmd11a | |
| **Psme3** | psme2 | | psme1 | | | | psme1, psme2 |
| **PTGDS** | ptgdsb.1 | | | | | | |
| **SDF2L1** | | | | sdf2l1 | | | |
| **SF3B4** | | sf3b6 | sf3a2 | | | | sf3a1, sfb1 |
| **SMARCA5** | | | smarce1 | | smarce1 | | |
| **SNRPD1** | | | snrpd1 | | snrpb, snrpd2 | | snrpd1, snrpd2 |
| | | | snrpd2 | | | | |
| **SNRPE** | | | | | | snrpe | |
| **SPC18** | spcs3 | | | | | | |
| **SRI** | | | | | | sri | |
| **TOMM34** | | | tomm20b | tomm5 | | | tomm20b |
| **Tpm3** | tpm1 | | | | tpma | | |
| **TSPAN-1** | tspan13a, tspan15 | tspan3a | | | tspan2a, tspan3a, tspan7b | | |
| **UBE2N** | ube2na | ube2e3 | ube2ib | ube2ia | ube2ib, ube2v1 | ube2ia | ube2v1 |
| **ZNF259** | | znf395b | ZNF276 | znf536 | znf395a, znf395b, znf593, znf609b | | |
